# Sox9 accelerates vascular ageing by regulating extracellular matrix composition and stiffness

**DOI:** 10.1101/2023.05.03.539285

**Authors:** Maria Faleeva, Sadia Ahmad, Steven Lynham, Gabriel Watson, Meredith Whitehead, Susan Cox, Catherine M Shanahan

**Affiliations:** BHF Centre of Research Excellence, School of Cardiovascular and Metabolic Medicine & Sciences, King’s College London, London, UK; Proteomics Facility, Centre of Excellence for Mass Spectrometry, King’s College London, UK; Randall Centre for Cell & Molecular Biophysics, Faculty of Life Sciences & Medicine, King’s College London, London, UK

**Author notes:** **Corresponding Author:** Professor Catherine M. Shanahan School of Cardiovascular and Metabolic Medicine & Sciences, King’s College London, London, SE5 9NU UK.

**Keywords:** Vascular Smooth Muscle Cells, Extracellular matrix, Extracellular vesicles, Sox9

## Abstract

**Rationale:** Vascular calcification and increased extracellular matrix (ECM) stiffness are hallmarks of vascular ageing. Sox9 (SRY-Box Transcription Factor 9) is a master regulator of chondrogenesis, also expressed in the vasculature, that has been implicated in vascular smooth muscle cell (VSMC) osteo-chondrogenic conversion.

**Objective:** Here, we investigated the relationship between vascular ageing, calcification and Sox9-driven ECM regulation in VSMCs.

**Methods and Results:** Immunohistochemistry in human aortic samples showed that Sox9 was not spatially associated with vascular calcification but correlated with the senescence marker p16. Analysis of Sox9 expression *in vitro* showed it was mechanosensitive with increased expression and nuclear translocation in senescent cells and on stiff matrices. Manipulation of Sox9 via overexpression and depletion, combined with atomic force microscopy (AFM) and proteomics, revealed that Sox9 regulates ECM stiffness and organisation by orchestrating changes in collagen expression and reducing VSMC contractility, leading to the formation of an ECM that mirrored that of senescent cells. These ECM changes promoted phenotypic modulation of VSMCs whereby senescent cells plated onto ECM synthesized from cells depleted of Sox9 returned to a proliferative state, while proliferating cells on a matrix produced by Sox9 expressing cells showed reduced proliferation and increased DNA damage, reiterating features of senescent cells. Procollagen-lysine, 2-oxoglutarate 5-dioxygenase 3 (LH3) was identified as a Sox9 target, and key regulator of ECM stiffness. LH3 is packaged into extracellular vesicles (EVs) and Sox9 promoted EV secretion, leading to increased LH3 deposition within the ECM.

**Conclusions:** These findings identify cellular senescence and Sox9 as a key regulators of ECM stiffness during VSMC ageing and highlight a crucial role for ECM structure and composition in regulating VSMC phenotype. We identify a positive feedback cycle whereby cellular senescence and increased ECM stiffening promote Sox9 expression which drives further ECM modifications that act to accelerate vascular stiffening and cellular senescence.

## Introduction

Vascular calcification is a life-threatening pathology that strongly associates with vascular ageing. It is caused by the abnormal deposition of calcium salts in the vessel intima and/or media and contributes to the development of atherosclerosis, as well as arterial stiffening(1). Vascular calcification is a cell-mediated process associated with modulation of vascular smooth muscle cells (VSMCs) to an osteo/chondrogenic phenotype characterised by loss of contractile markers (2,3), increased expression of key transcription factors (TFs) associated with developmental osteo/chondrogenesis (4,5)and increased extracellular vesicle (EV) release(6,7).

The transition of VSMCs to an osteo/chondrogenic phenotype has been tightly linked with cellular ageing. Senescent VSMCs are increased in the calcified vessel wall and calcify more readily *in vitro* (8–10). Senescent cells are also pro-inflammatory, expressing a senescence-associated secretory phenotype (SASP), and secreting a large volume of cytokines, chemokines and interleukins, as well as EVs (11). Secreted EVs are involved in both physiological and pathological processes, mediating intracellular communication but also depositing in the extracellular matrix (ECM) to form a nidus for mineralization(12). During senescence, there are changes in cargo loading of EVs, resulting in an increased abundance of calcification-promoting factors. Furthermore, these EVs have the potential to modulate the phenotype of neighbouring cells (13).

In addition to mineralisation, the ageing vasculature also undergoes dramatic changes in ECM composition and structure that contribute to vascular stiffening. In the aorta there is a decrease in elastin deposition and an increase in its fragmentation, increased collagen deposition and an increase in the crosslinking of collagen fibres, with some of these changes also implicated in promoting calcification (14,15). Many of these changes to the ECM have been attributed to ‘wear and tear’ or oxidation processes or have been studied in the context of disease. Few studies have considered how VSMC ageing and senescence may act to directly modulate ECM ageing, or to establish the factors that might regulate this interplay. Indeed, ECM proteins play a dual role in regulating both the integrity of the vasculature and extracellular cell signalling yet, how ECM ageing impacts on VSMC phenotype remains poorly understood.

RUNX Family Transcription Factor 2 (Runx2) and SRY-Box Transcription Factor 9 (Sox9) are master regulators of bone and cartilage differentiation. These TFs cooperate to orchestrate expression of numerous matrisomal proteins required to form and promote, or not, the mineralisation of these specialist tissues (16). Both are also expressed in the calcified vasculature (5). However, while the role of Runx2 in activating the expression of osteogenic genes during vascular calcification and ageing has been well characterized (17–20), less is known about the expression and role of Sox9. Interestingly, Sox9 plays a key role during vascular development. In sclerotomal progenitor cells Sox9 is vital for cell fate specification between VSMCs and chondrocytes and must be silenced via Notch signalling (21) to enable VSMC differentiation. This silencing allows de-repression of myocardin, a key TF regulating expression of smooth muscle marker genes such as SM22-α and α-SMA. Sox9 also induces VSMC de-differentiation *in vitro* by decreasing expression of contractile genes (22–24). On the other hand, maintenance of Sox9 expression during development facilitates the expression of numerous ECM genes, such as Collagen2, 9, and 11, essential for chondrocyte differentiation (25,26). Vascular injury and ageing can prompt a decrease in Notch signalling (27) and potentially induce re-expression of Sox9 (21,23). Consistent with this notion, expression of Sox9 had been demonstrated in mouse models of arteriosclerosis where it has been shown to play roles in ECM-reorganization through its upregulation of Collagen 2 (23) as well as in calcification (23), via the regulation of proteoglycan 4 (PRG4) (28). In the human vasculature Sox9 expression has been shown to occur in aged and calcified aortic tissue (29) however, its role in ECM remodelling and calcification has been largely overlooked.

In this study we describe a novel role for Sox9 in human vascular ageing. We show that Sox9 expression in the vasculature correlates with VSMC senescence and demonstrate that Sox9 is mechanosensitive in aged VSMCs. We also show that it is a critical mediator of ECM stiffness through its activation of the collagen modifier LH3 and its role in driving increased secretion of LH3 in EVs. Importantly the ECM produced in response to Sox9 mimics key features of the senescent ECM and acts as a potent inhibitor of VSMC proliferation while concomitantly driving DNA damage and inflammation to accelerate VSMC ageing.

## Results

### Sox9 expression in the vessel media correlates with loss of contractile markers and ageing but not calcification

We performed immunohistochemistry (IHC) on human aortic tissue samples from young and old patients with and without vascular calcification. Nuclear Sox9 expression was detectable in the vessel media predominantly in VSMCs (Fig 1a). As expected, there was a strong correlation between patient age and calcification, and age and expression of p16 a marker of cell senescence (S Fig1a-c). However, we found no association between Sox9 expression and calcification (Fig S 1 d,e). We observed some non-calcified areas with high levels of Sox9, whilst areas of high calcification were negative for Sox9 staining (S Fig 1a). Using an *in vitro* calcification assay we verified that calcification propensity was not increased in Sox9 overexpressing VSMCs suggesting there is not a direct relationship between Sox9 and mineralisation in human cells (Fig S1f,g). However, we did find a correlation between Sox9 and the senescence marker p16. Elevated expression of Sox9 was also associated with decreased α-SMA expression (Fig 1b,c) with p16 also showing the same negative correlation with α−SMA (Fig 1d). IHC of Sox9 and p16 in serial sections revealed more p16 positive cells than Sox9 positive cells in any given section, with the majority of Sox9 positive nuclei also staining positive for p16, typically in areas that were cell poor and matrix-rich (Fig 1e,f). This suggested that senescent VSMCs may be providing an environment that promotes elevated Sox9 expression.

**Figure 1:**
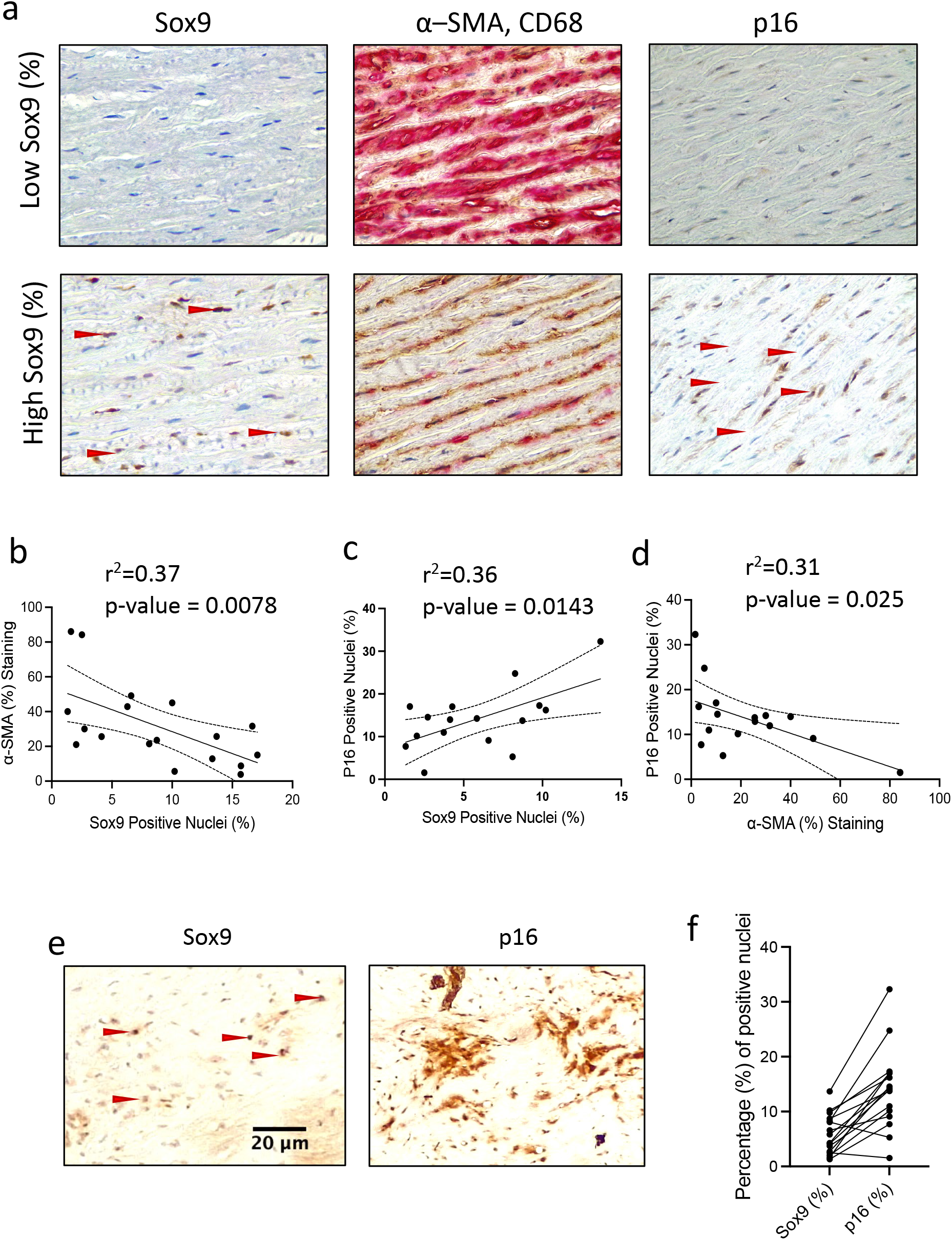
Sox9 expression in VSMCs correlates with increased cellular senescence and decreased expression of α-smooth muscle actin (α-SMA). (a) Sox9, α-SMA, and CD68 and p16 staining in the medial layer of aorta from human subjects. (b) Correlation of Sox9 positive nuclei with a-SMA and (c) p16 positive nuclei. Correlation of α-SMA with (d) p16 positive nuclei. Red arrows highlight positively stained nuclei for Sox9 and p16, n=16. Correlation was performed using a pearson test. (e) IHC of serial sections stained for Sox9 and p16 within an identical area of the media (f) Bar graph showing percentages of Sox9 and p16 nuclei in each sample.

### Sox9 is mechanosensitive during VSMC senescence

The association between p16 and Sox9 *in vivo* led us to investigate whether Sox9 is increased during cellular senescence. We cultured primary human VSMCs to replicative senescence using serial passaging and found that senescent VSMCs showed increased expression of Sox9 at the RNA level, but not consistently at the protein level (Fig2a-c). Immunofluorescence (IF) however showed more nuclear expression of Sox9 in senescent VSMCs compared to their early passage counterparts (Fig 2d-e). To investigate whether Sox9 can promote VSMC senescence, Sox9 was overexpressed in early passage (hereafter termed ‘young’) VSMCs using adenoviral transduction (Sox9 OE). This resulted in a decrease in contractile markers such as α-SMA consistent with its role in regulating VSMC differentiation (S Fig2a-c). However, minimal changes in p16 gene expression were observed. Sox9 depletion (Sox9 KO) in senescent cells also had no effect on p16 (S Fig 2d-f) suggesting other factors are involved in Sox9 regulation during VSMC ageing.

**Figure 2:**
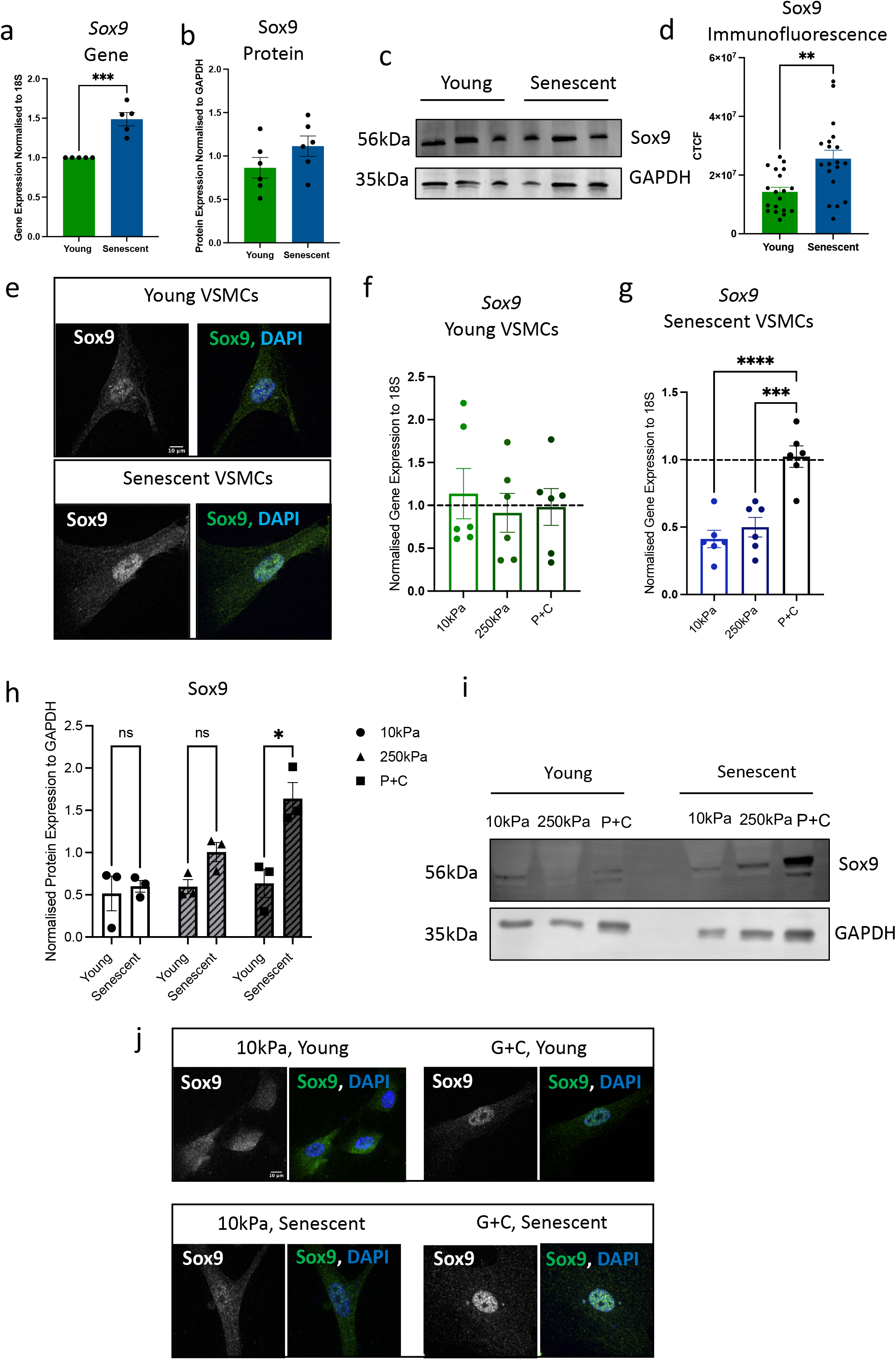
Sox9 is mechanosensitive in senescent VSMCs. (a,b,c) Sox9 gene and protein expression in young and senescent VSMCs, quantified by RT-qPCR and Western blotting, n=6. (d,e) Quantification of Corrected Total Cell Fluorescence (CTCF) and immunofluorescence of Sox9 in young and senescent VSMCs, n=18. Unpaired Student’s t test, **p-value<0.01, ***p<0.005. Sox9 (f,g) Gene and (h,i) protein expression, quantified by RT-qPCR and western blotting, of young and senescent VSMCs plated on hydrogels of 10kPa, 250kPa stiffness, and plastic coated with collagen (P+C), n=6, n=3, respectively. One-way ANOVA with Tukey post hoc test, *p-value<0.05, ***p<0.005, ****p<0.0001. (j) Immunofluorescence of Sox9 when plated on hydrogels of 10kPa stiffness and glass coated with collagen (G+C) in young and senescent VSMCs. Sox9 in grey and green, nuclear DAPI staining in blue.

In chondrocytes Sox9 is mechanosensitive and shows increased expression in stiff environments (30). As vascular ageing is associated with increased ECM stiffness, we tested whether Sox9 might be mechanosensitive in VSMCs. Young and senescent VSMCs were plated onto hydrogels of different stiffness (Fig 2f-j). While young VSMCs showed no changes in Sox9 gene expression, senescent VSMCs showed markedly increased expression on stiff collagen coated plastic at both the gene and protein level (Fig2 f-i). Moreover, IF showed that when plated on soft matrices, Sox9 is diffusely localized in the cytoplasm, and this was most pronounced in young cells with senescent cells already showing nuclear localisation. When plated on stiff matrices, Sox9 translocated into the nucleus in both young and senescent cells, indicating a potential increase in its transcriptional activity (Fig 2j). This was confirmed using RT-qPCR for Sox5 and Sox6, two transcriptional targets of Sox9, that also showed increased expression on stiff surfaces (S Fig2 g-j).

### Sox9 regulates VSMC extracellular matrix organisation and stiffness

Sox9 has been shown to regulate an array of extracellular matrix (ECM) related genes in chondrocytes (26) and mouse VSMCs (23). Thus, we next wondered whether Sox9 may regulate ECM changes during vascular ageing. To examine the effects of ageing on matrix stiffness, ECM was synthesized from young and senescent VSMCs, the cells removed, and the stiffness of the native ECM measured by atomic force microscopy (AFM). This revealed an increase in stiffness of ECM produced by senescent VSMCs compared to that synthesized by young cells (Fig 3a). We next analysed the ECM by IF (Fig 3b). Fibronectin staining revealed that ECM synthesized by young cells showed an organised fibrillar pattern. In contrast the senescent ECM showed a woven, matted appearance with disorganized fibers. Alignment by Fourier Transform (AFT) analysis confirmed quantitative differences in matrix alignment. Mapping the topography of the fluorescent signal of fibronectin also revealed that the senescent ECM adopts a flat appearance compared to the peaks and valleys seen in the young ECM.

**Figure 3:**
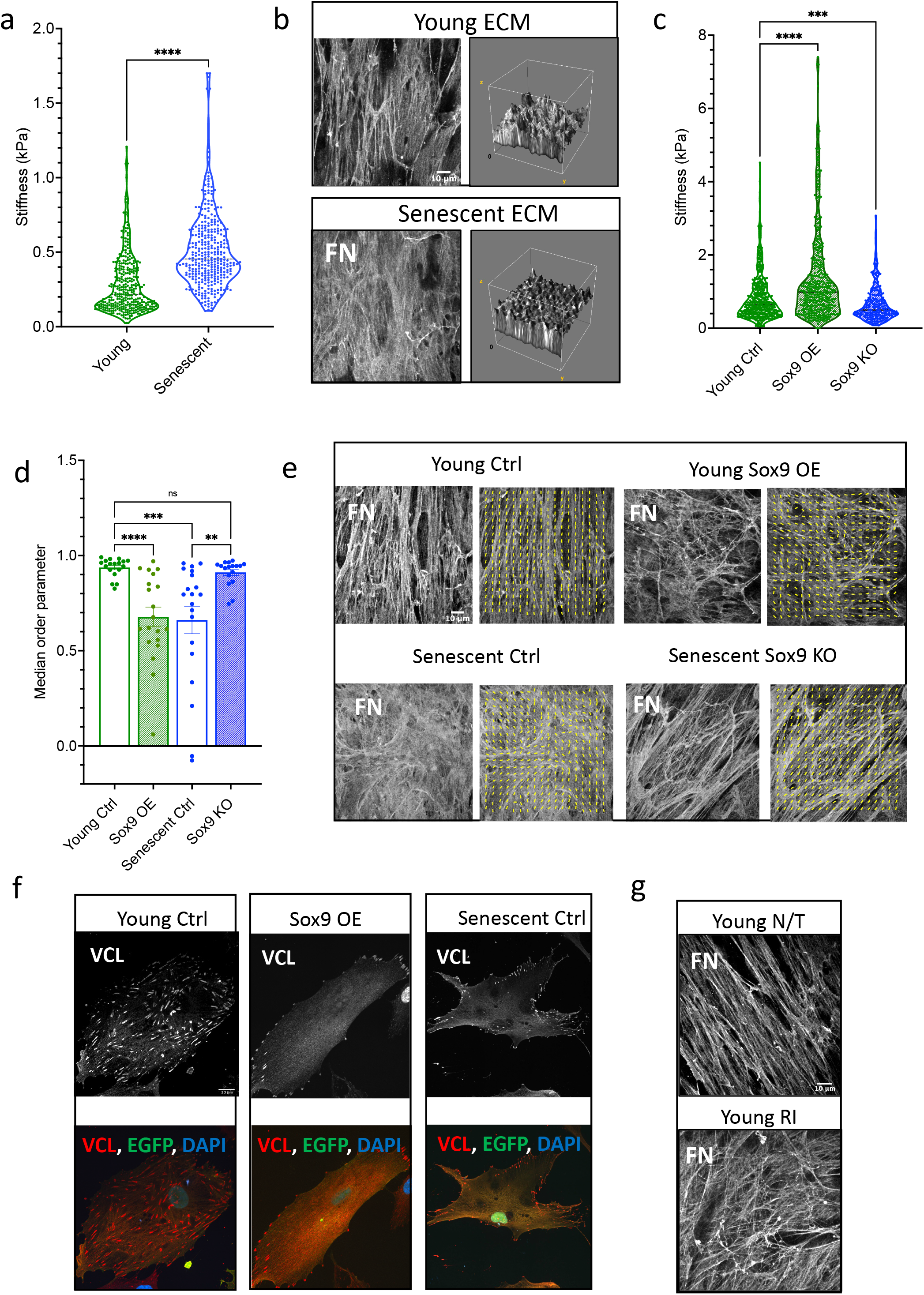
VSMC senescence and Sox9 expression regulate the ECM. (a) Atomic force microscopy (AFM) measurements of ECM stiffness synthesized from young and senescent VSMCs. Normality rejected via Shapiro-Wilk test, Unpaired Mann-Whitney test, ****p-value<0.0001. Stiffness measurements taken from ECM synthesized from three different donors; 35F, 38F, 33F. (b) Representative fibronectin immunofluorescence staining of young and senescent ECM. Topology of fibronectin staining presented to the right-hand side of the immunofluorescence image. (c) AFM measurements of ECM synthesized from young VSMC transfected with EGFP (Young Ctrl), Sox9 overexpression adenovirus (Sox9 OE), and Sox9 adenoviral knockout (Sox9 KO). Normality rejected via Shapiro-Wilk test, Unpaired Mann-Whitney test, n=20 ***p-value<0.005, ****p-value<0.0001. (d) Median order parameter of analyzed fibronectin fibre alignment in ECM synthesized from Young Ctrl, Sox9 OE, Senescent Ctrl, and Sox9 KO VSMCs. (e) Representative immunofluorescence staining of fibronectin in ECM synthesized from Young Ctrl, Young Sox9 OE, Senescent VSMCs transduced with EGFP (Senescent Ctrl), and Senescent cells with Sox9 knockout (Sox9 KO), with fiber alignment indicated in yellow lines. (f) Representative immunofluorescent images of vinculin (VCL) in Young Ctrl, Sox9 OE, and Senescent Ctrl. VCL in grey/red, EGFP in green, nuclear staining (DAPI) in blue. (g) Representative immunofluorescence staining of fibronectin in ECM synthesized from young VSMCs with no treatment (Young N/T) and young VSMCs treated with Rock Inhibitor (Young RI).

To investigate the potential role of Sox9 in regulating ECM stiffness and fibre organisation, we overexpressed or depleted Sox9 in young and senescent VSMCs respectively and analysed the resulting ECM (Fig 3). We found that Sox9 overexpression led to an increase in ECM stiffness and a decrease in parallel fibre alignment, creating an ECM resembling that produced by senescent VSMCs. Conversely, its depletion in senescent cells resulted in decreased stiffness and an increase in parallel fibres quantitatively no different from the alignment observed in young ECM (Fig 3c-e). To understand the mechanism behind this, we tested whether Sox9 regulates fibre alignment through focal adhesions (FA), which anchor cells to the ECM. Overexpression of Sox9 resulted in a reduction and redistribution of vinculin, a marker for FAs (31). FAs were present over the entire cell area in young control cells but only present around the perimeter of the cell membrane in response to Sox9. This was similar to the FA localisation observed and previously described in senescent VSMCs (Fig 3f). Importantly, we could replicate the disorganised fibre alignment found in senescent or Sox9 expressing VSMCs by treating young VSMCs with a ROCK inhibitor to reduce cellular contractility to mimic the effect of Sox9 on reducing VSMC contraction (Fig 3g). These findings suggest that Sox9 plays a crucial role in regulating ECM stiffness and fibre organisation in part via regulation of cellular contractility and FA localisation.

### Sox9 Regulated ECM regulates VSMC Phenotype

We next investigated whether the Sox9-modulated ECM could influence VSMC phenotype and senescence. To do this, we again synthesized ECM from young and senescent VSMCs with Sox9 overexpression or depletion, respectively. We then replated fresh young or senescent VSMCs on top of their corresponding decellularized ECM (Fig 4a). We found that young VSMCs plated on ECM synthesized from young VSMCs overexpressing Sox9 had increased expression of p16 and p21, as well as elevated IL6, an inflammatory SASP marker (32), compared to cells plated on EGFP control young ECM. In contrast, senescent cells plated on matrices synthesised from senescent VSMCs depleted of Sox9 exhibited a downregulation of both p16 and p21, as well as IL6 (Fig 4b-c). Importantly, cell proliferation, measured via Edu incorporation, and DNA damage analysis were consistent with the changes in expression of these senescent cell cycle and inflammatory markers observed in the cells plated on Sox9 modified matrices. Young VSMCs plated on young Sox9 overexpressing ECM had reduced cell proliferation and an increased number of DNA double-strand breaks shown by γH2Ax foci consistent with DNA damage driving cell cycle arrest and senescence. Conversely, senescent cells plated on senescent Sox9 knockout ECM were able to re-enter the cell cycle as shown by increased Edu incorporation without any change in DNA damage (Fig 4 d-f). Furthermore, we observed that the orientation of the VSMCs was highly dependent on the ECM. Both young and senescent VSMCs when plated on young control and senescent Sox9 knockout matrices exhibited parallel orientation to one another (Fig4g,h). VSMCs plated on young Sox9 overexpression and senescent control ECM exhibited random orientation, frequently aligning at perpendicular angles with each other (Fig 4g,h). This change in cell orientation is likely due to a lack of fibronectin fibre definition and alignment as shown previously in Fig 3. To rule out that the changes in gene expression observed were due to changes in matrix stiffness we performed RT-PCR for p16, p21 and IL6 on hydrogels of different stiffness. None were mechanosensitive (S Fig 3a-f) while expression of the mechanosensitive proteins, α-SMA and SM22-a, which are reduced on soft matrices did not show any changes in response to Sox9 mediated ECM modulation (S Fig 3i-l). This suggests that it is ECM composition, not stiffness or alignment, that is driving these changes in gene expression.

**Figure 4:**
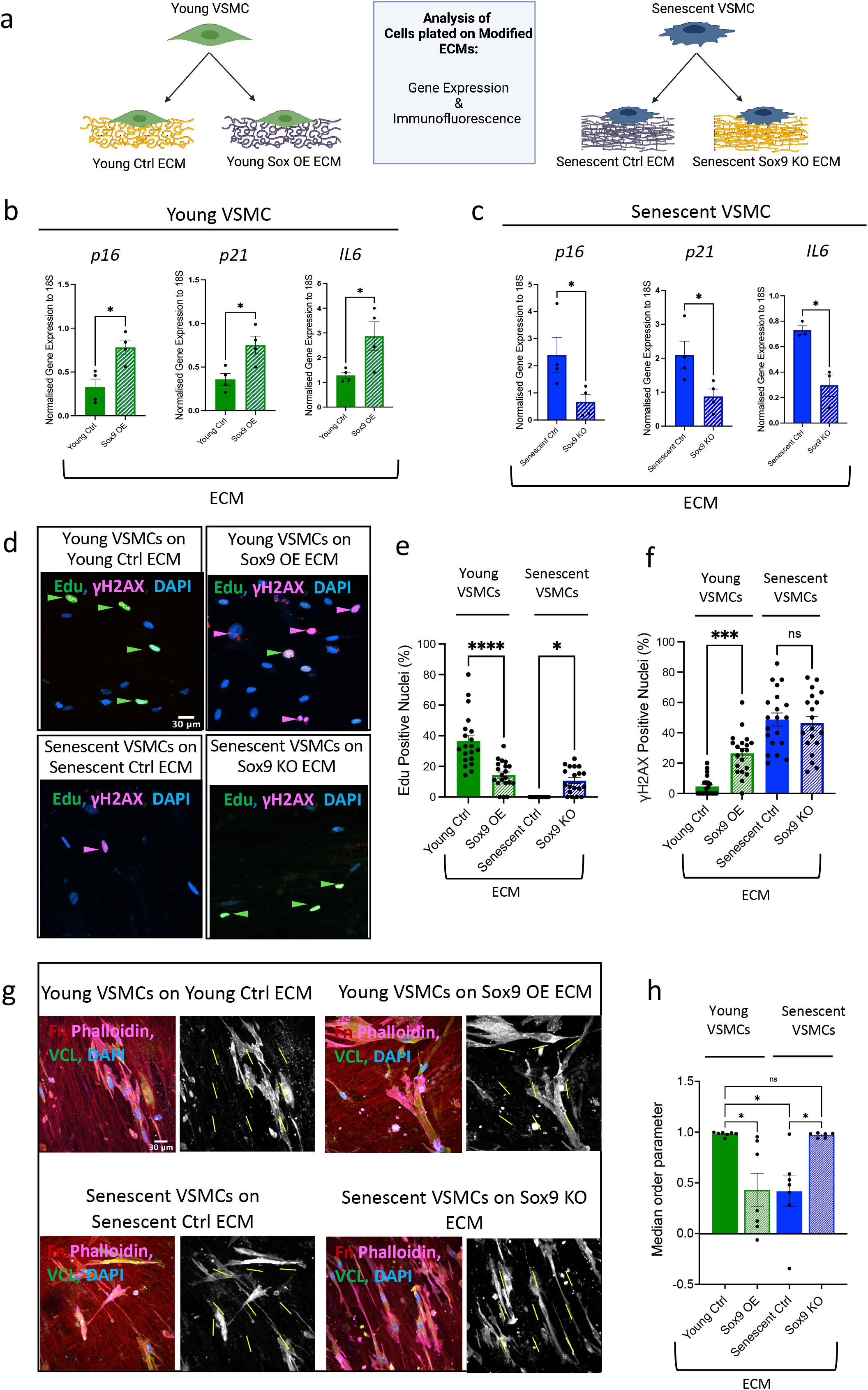
Composition of Sox9-regulated ECM regulates VSMC phenotype. (a) Schematic of experimental protocol. Young VSMCs were plated on decellularized ECM originally synthesized from young VSMCs transduced either with EGFP (Young Ctrl) or Sox9 overexpression adenovirus (Sox9 OE). Senescent VSMCs were plated on decellularized ECM originally synthesized from senescent VSMCs transduced either with shEGFP (Senescent Ctrl) or Sox9 knockout adenovirus (Sox9 KO). Gene expression, quantified by RT-qPCR, of *p16*, *p21*, and *IL6* from (b) young and (c) senescent VSMCs plated on transduced ECMs. N=4, Unpaired Student’s t-test, *p-value<0.05. (d) Representative IF and quantification (%) of (e) Edu and (f) *γ*H2AX when VSMCs plated on transduced ECMs. Edu, γH2AX, and nuclear DAPI staining in green, magenta, and blue, respectively, n=20. One-way ANOVA with Tukey post hoc test, *p-value<0.05, ***p-value<0.005, ****p-value<0.0001. (g) Representative IF staining of young and senescent VSMCs when plated on transduced ECMs, with fibronectin (Fn) in red, Phalloidin in magenta, Vinculin (VCL) in green, and nuclear staining (DAPI) in blue. Adjacent black and white images of vinculin staining with yellow lines indicating cellular alignment within the frame. (h) quantified median order parameter of cellular alignment of young and senescent VSMCs plated on ECM synthesized from Young Ctrl, Sox9 OE, Senescent Ctrl, and Sox9 KO VSMCs. One-way ANOVA with Multiple Comparisons, n=6, *p-value<0.05.

### Sox9 Regulates Protein Ratios Deposited in the ECM

To gain insights into the specific components of the ECM that are directly regulated by Sox9, we next conducted Mass spectrometry analysis on young and senescent decellularized matrix with and without Sox9 overexpression and Sox9 knockout, respectively (Fig 5). Principle component analysis (PCA) and differentially expressed protein (DEP) clustering showed young and senescent ECM formed distinct clusters (S. Fig 4a). Notably, ECM from young Sox9 overexpression clustered most closely with senescent control ECM, while senescent Sox9 knockout ECM formed a separate cluster more closely related to young control ECM, as illustrated by heatmap (Fig5a). GO analysis firstly comparing young and senescent control ECM showed the main pathways downregulated were focal adhesion, basement membrane and ECM while upregulated pathways included extracellular exosome (S Fig 4b). Remarkably, we found similar pathway changes when Sox9 overexpression ECM was compared with young control ECM with a decrease in ECM-related proteins and an increase in proteins associated with extracellular exosomes and cytoplasmic proteins. The opposite was observed when senescent Sox9 knockout ECM was compared with senescent ECM which showed an increase in ECM related proteins and a decrease in exosome and cytoplasmic components (Fig5b). Further comparative analysis to determine if the proteins changing within the ECM were similar showed there was a 50% overlap in the differentially expressed proteins between senescent control and young Sox9 overexpression ECM when both were compared independently with the young control ECM. The majority of these proteins were glycoproteins and collagens in the matrisomal protein group (Fig5c). Volcano plots highlighted Col15a1 as having the highest negative log fold change in both the senescent control and in the young Sox9 overexpression ECM as compared to the young control ECM (Fig 5d). Other matrisomal proteins that were reciprocally changed in each group, and also differentially expressed between young and senescent control ECM, included collagen 4 and Elastin microfibril interfacer 1 (EMILIN1). The senescent Sox9 knockout ECM also exhibited a strong upregulation of Lysyl Oxidase Like 4 (LOXL4) a key modulator of collagen and elastin cross-links (Fig 5d). There were also reciprocal changes in a number of Annexin (Anx) proteins, which are abundant EV cargoes (33).

**Figure 5:**
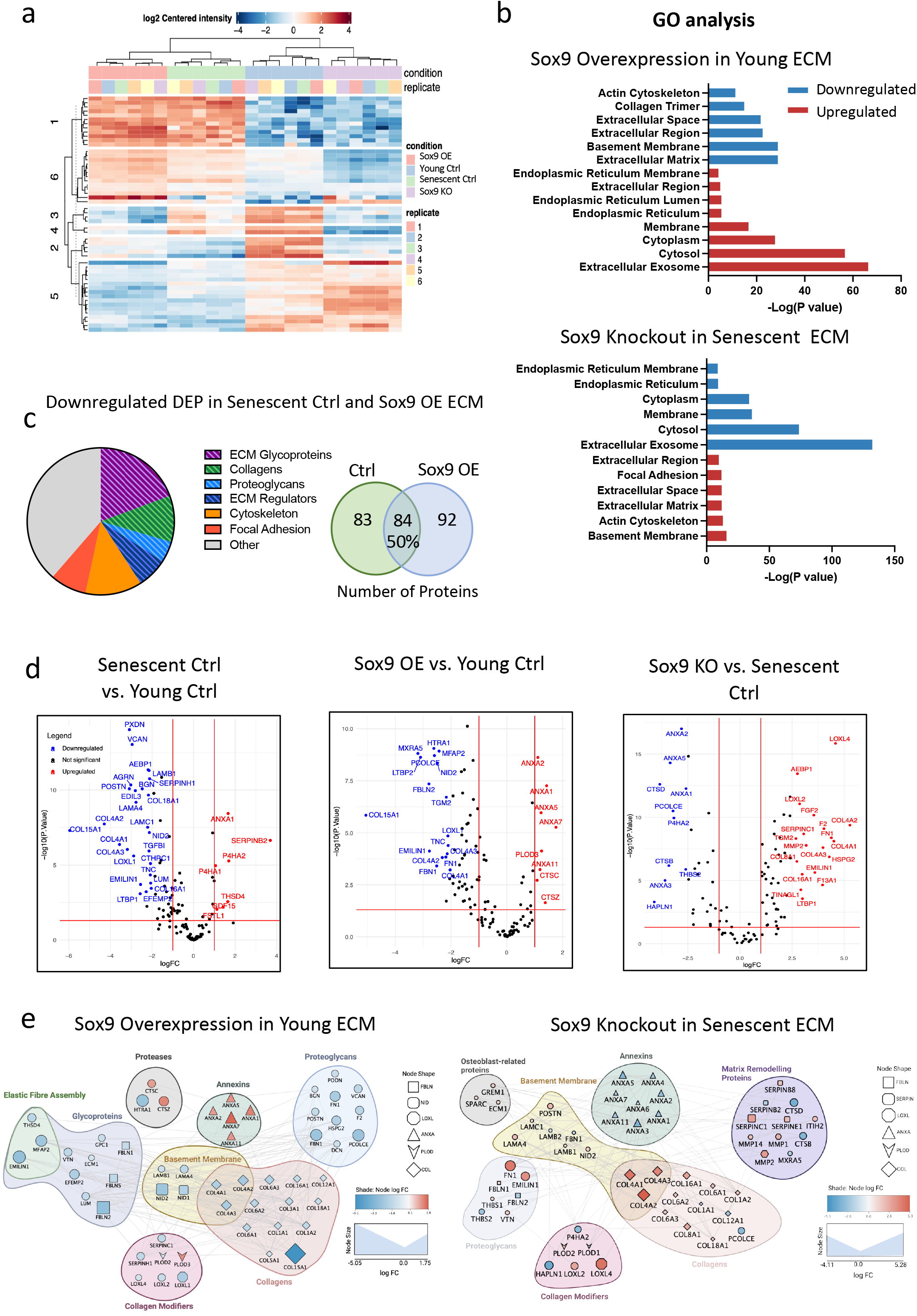
Sox9 drives the ECM composition towards a senescent-like phenotype (n= 6 injections for each condition from 35-year-old female (35F) and 38-year-old female (38F)). (a) Heatmap of clustered differentially expressed proteins (DEP), showing the similarly of protein expression between ECM synthesized from young VSMCs treated with control EGFP adenovirus (Young Ctrl) and senescent VSMCs with Sox9 knockout adenovirus (Sox9 KO), and between young VSMCs transduced with Sox9 overexpression adenovirus (Sox9 OE) and senescent VSMCs transduced with control shEGFP adenovirus (senescent Ctrl). (b) Gene ontology (GO) analysis of differentially expressed proteins (DEP) in Sox9 OE ECM as compared to Young Ctrl, and Sox9 KO as compared to Senescent Ctrl. (c) Pie chart and Venn-diagram of downregulated DEP both in the Sox9 OE and Senescent Ctrl ECM. (d) Volcano plots of DEP when comparing Senescent Ctrl against Young Ctrl, Sox9 OE against Young Ctrl, and Sox9 KO against Senescent Ctrl. (e) Protein-protein interaction network showing interactions between significantly upregulated and downregulated proteins in Sox9 OE ECM as compared to the Young Ctrl ECM, and in Sox9 KO ECM as compared to the Senescent Ctrl ECM.

To further understand the functional relationships between the differentially expressed matrisomal proteins regulated by Sox9 we constructed protein-protein interaction (PPI) networks. This revealed that the most downregulated proteins in the young Sox9 expression ECM were glycoproteins and collagens. Amongst these were a group of proteins associated with elastic fibre assembly and deposition including EMILIN1 and Fibulins1/2 (FBLN1/2). There was also a clear subset of basement membrane components such as Collagens 4a1-2 (Col4a1-2) and 6 (Col6a1-2) and Nidogens 1/2(NID1/2). Collagen modifier proteins including LOXL2/4 were mostly downregulated however two modifiers, Cathepsin C (CTSC) and Procollagen-lysine,2-oxoglutarate 5-dioxygenase 3 (*Plod3*/LH3) were upregulated. A large subset of annexin proteins including those enriched in VSMC EVs were also upregulated (33). Conversely, ECM synthesized from senescent VSMCs depleted of Sox9 showed the opposite changes. There was clear upregulation of the same elastin modifiers as well as collagens and basement membrane proteins. ECM remodelling proteins were also upregulated including LOXL4 (Figure 5e). Interestingly, both PLOD1/2 were downregulated as were annexins. We did not see any changes in key chondrocyte Sox9 targets such as collagen 2 and 11. We validated the changes in basement membrane components using RT-PCR and IF which revealed clear defects in basement membrane assembly of Col 4 in both senescent and young Sox9 expressing VSMCs further highlighting the key role Sox9 plays in regulating these protein networks. (S Fig 4c-f).

### Sox9 regulates the posttranslational modification of collagen fibrils via LH3

Most interestingly, we observed that Sox9 upregulated collagen modifiers within the ECM that could potentially be responsible for the changes in ECM stiffness. *Plod3*/LH3, a protein with dual cross-linking and glycating properties, stood out as one of the few proteins that was upregulated in the young Sox9 overexpression ECM (Fig 6). To validate the changes in LH3 in response to Sox9, we overexpressed and depleted Sox9 in young and senescent VSMCs and examined LH3 expression using RT-PCR and Western blot. Results showed that *Plod3* was upregulated in young Sox9 overexpressing VSMCs and downregulated in senescent Sox9 knockout VSMCs as compared to the young and senescent controls, respectively (Fig 6a). On the protein level, LH3 was upregulated with Sox OE, but not downregulated in Sox9 KO as compared to their respective controls (Fig 6b,c). IF showed that LH3 was consistently localised to the Golgi apparatus with no change in localisation either with senescence or Sox9 expression (S Fig 5a). However, examination of the decellularized ECM using IF showed that LH3 deposition was markedly increased in the young Sox9 overexpression and senescent control conditions compared with the young control and senescent Sox9 knockout ECM where it was decreased (Fig 6 d-g). LH3 also showed a distinctive pattern change in the Sox9 overexpression which mirrored the pattern seen in the senescent control ECM, adopting an increased fibrillar deposition pattern (Fig 6g). This suggested that increased deposition of LH3 into the matrix, rather than increased intracellular protein synthesis, maybe involved in ECM modulation.

**Figure 6:**
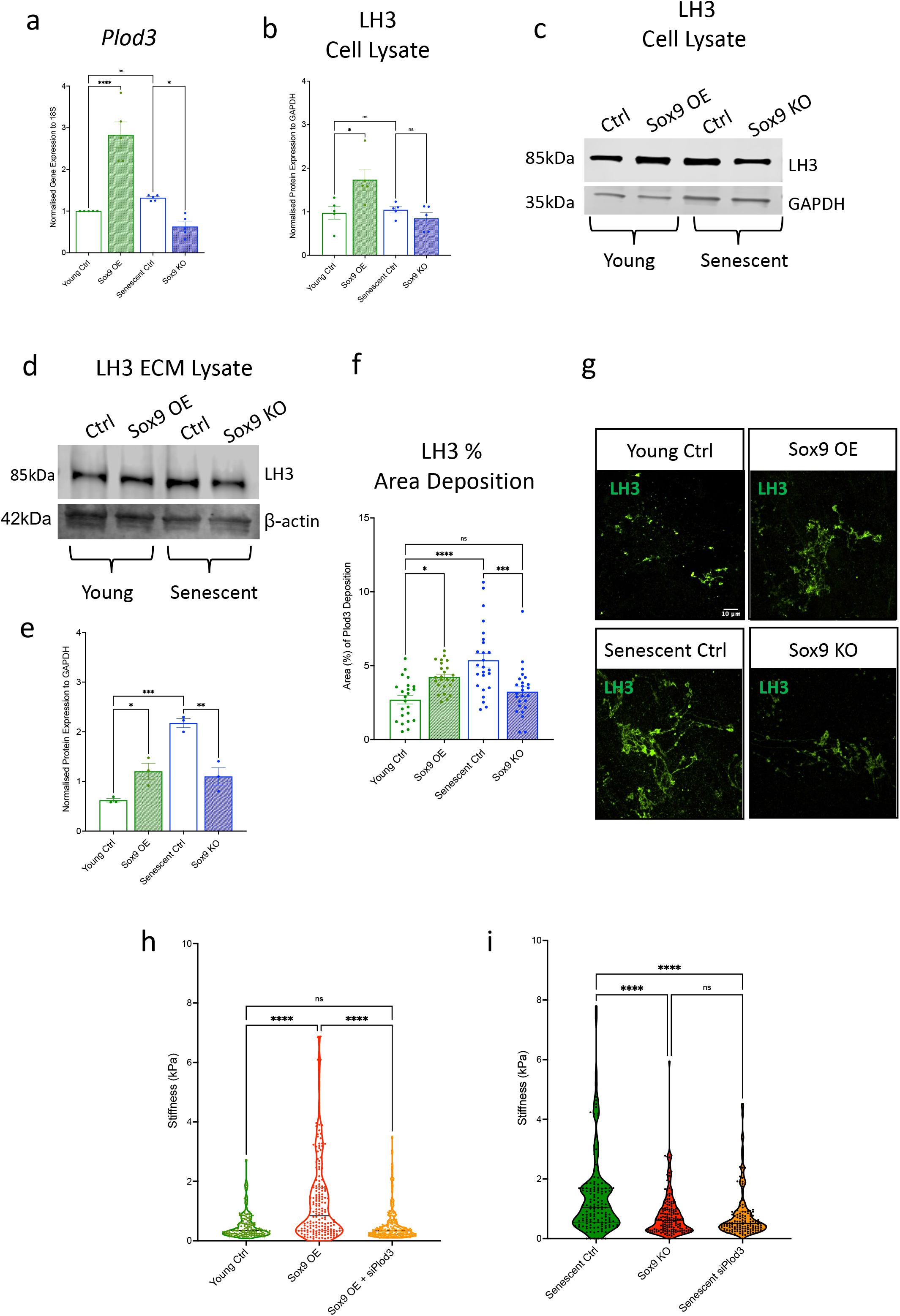
*Plod3*/LH3 in VSMCs expression and deposition into the ECM is regulated by Sox9 expression. (a) Gene expression, quantified via RT-qPCR, of *Plod3* in young VSMCs transduced with EGFP control adenovirus (Young Ctrl), Sox9 overexpression adenovirus (Sox9 OE), and senescent VSMCs transduced with shEGFP control adenovirus (Senescent Ctrl) and Sox9 knockout adenovirus (Sox9 KO), n=5. Quantification and representative Western blot of LH3 protein expression in Young Ctrl, Sox9 OE, Senescent Ctrl, and Sox9 KO VSMCs (b,c) cell lysate, n=5, and (d,e) ECM, n=3. (f,g) Quantification and representative immunofluorescence of fluorescent LH3 signal when deposited into the ECM synthesized from Young Ctrl, Sox9 OE, Senescent Ctrl, and Sox9 KO VSMCs, n= 25. One-way ANOVA with Tukey post hoc test, *p-value<0.05, ***p-value<0.005, ****p-value<0.0001. (h) Atomic force microscopy (AFM) stiffness measurements (kPa) in decellularized ECM synthesized from (e) Young Ctrl, Sox9 Sox9 OE, and Sox9 OE with Plod3 knockout (siPlod3) and (i) Senescent Ctrl, Sox9 KO, and Plod3 knockout (Senescent siPlod3). Measurements were taken from ECM synthesized from three isolates, 35F, 38F, and 33F. Shapiro-Wilk test for normality distribution was rejected. One-way ANOVA Kruskal-Wallis test, ****p-value <0.0001.

We next tested whether LH3 plays a role in Sox9-driven ECM stiffness by depleting *Plod3* using RNAi in young Sox9 overexpressing and in senescent control VSMCs (S Fig 5b). AFM stiffness measurements revealed a marked reduction of ECM stiffness in matrices that were synthesised from young VSMCs treated with Sox9 overexpression adenovirus in combination with siPlod3 as compared to the young Sox9 overexpression ECM. There was also no difference in stiffness when compared to the young control ECM (Fig 6h). Knocking out *Plod3* in senescent VSMCs resulted in a reduction of ECM stiffness as compared to the senescent control to a stiffness equivalent to that observed in Sox9 knockout ECM (Fig 6i). This demonstrated that LH3 may be a primary target driving Sox9-dependent ECM stiffness in ageing.

To validate our findings *in vivo*, we again conducted IHC on aortic patient tissue samples testing for LH3, Sox9 and p16 (Fig 7). We found that LH3 was increased in the aorta of aged individuals and that there was a correlation between elevated expression of LH3 and Sox9 as well as LH3 and p16 mirroring the previous correlations observed for Sox9 and p16 in patient tissue samples (Fig 7a-c).

**Figure 7:**
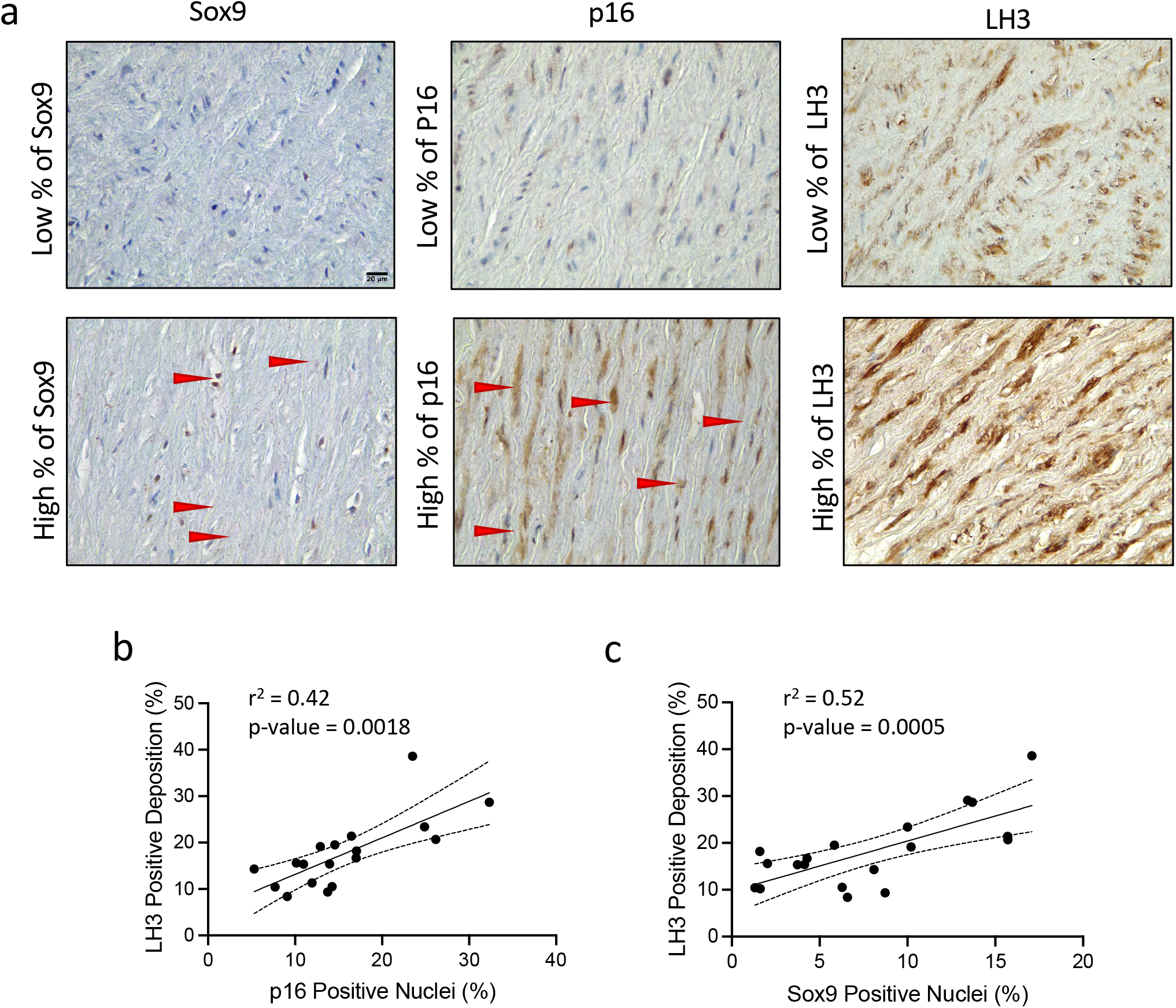
LH3 deposition increases in the medial aortic layer with age and Sox9 expression and regulates extracellular matrix (ECM) stiffness. (a) Immunohistochemistry of Sox9, p16, and LH3 staining in the aortic medial layer. The red arrows highlight positive staining of Sox9 and p16. Positive staining of p16, Sox9, and LH3 in brown. Correlation of LH3 positive staining (%) with (b) p16 and (c) Sox9 Positive nuclei. Correlation determined by Spearman’s correlation, n=16.

### LH3 deposition into the ECM is regulated via extracellular vesicles

EVs carry cargo from the cytosol that deposit within the extracellular space (34). The proteomics dataset showed that the upregulated proteins deposited in the ECM with Sox9 overexpression and with senescence were related to EVs (Fig 8a). We again performed comparative analysis to determine what proteins were differentially expressed between senescent control and young Sox9 overexpression ECM when both were compared independently with the young control ECM. This showed 171 proteins were in common representing a 61% overlap between the groups. Sub-classifying these in common upregulated proteins revealed that several were related to the regulation of intracellular transport, such as early endosome antigen 1 (EEA1) and Ras-related protein Rab21 (Rab21)(Fig 8b). As LH3 has been shown to be trafficked into the ECM in Collagen IV carriers (35), we hypothesized that EVs may drive the increased deposition of LH3 into the ECM in VSMC senescence. To validate changes in EV secretion and deposition in response to Sox9 overexpression and VSMC senescence we performed a CD63 bead capture assay and IF for the EV marker CD63 to quantify exosome secretion (Fig 8c) and deposition into the ECM respectively (Fig 8d,e). These assays showed that Sox9 overexpression, as well as VSMC senescence, increased EV secretion and deposition, while Sox9 knockout resulted in the opposite effect, decreasing EV secretion and deposition. Using differential ultracentrifugation to isolate different EV populations, we showed that LH3 was detectable in all EV subtypes including apoptotic bodies (Abs), micro-vesicles (miEV) and small vesicles (sEVs) in both young and senescent VSMCs (Fig 8f), with a trend to increased LH3 loading into sEVs in senescent cells (S Fig 6). To investigate the role of sEVs in the secretion of LH3, we inhibited EV secretion using 3-O-methylsphingomyelin (3-OMS) in both young and senescent VSMCs during ECM synthesis. We observed a reduction in LH3 deposition into the ECM, further validating its trafficking into the ECM via EVs. These findings support the notion that Sox9 drives increased release of EVs into the ECM during senescence, ultimately leading to the enhanced deposition of LH3 and the modulation of ECM mechanics.

**Figure 8:**
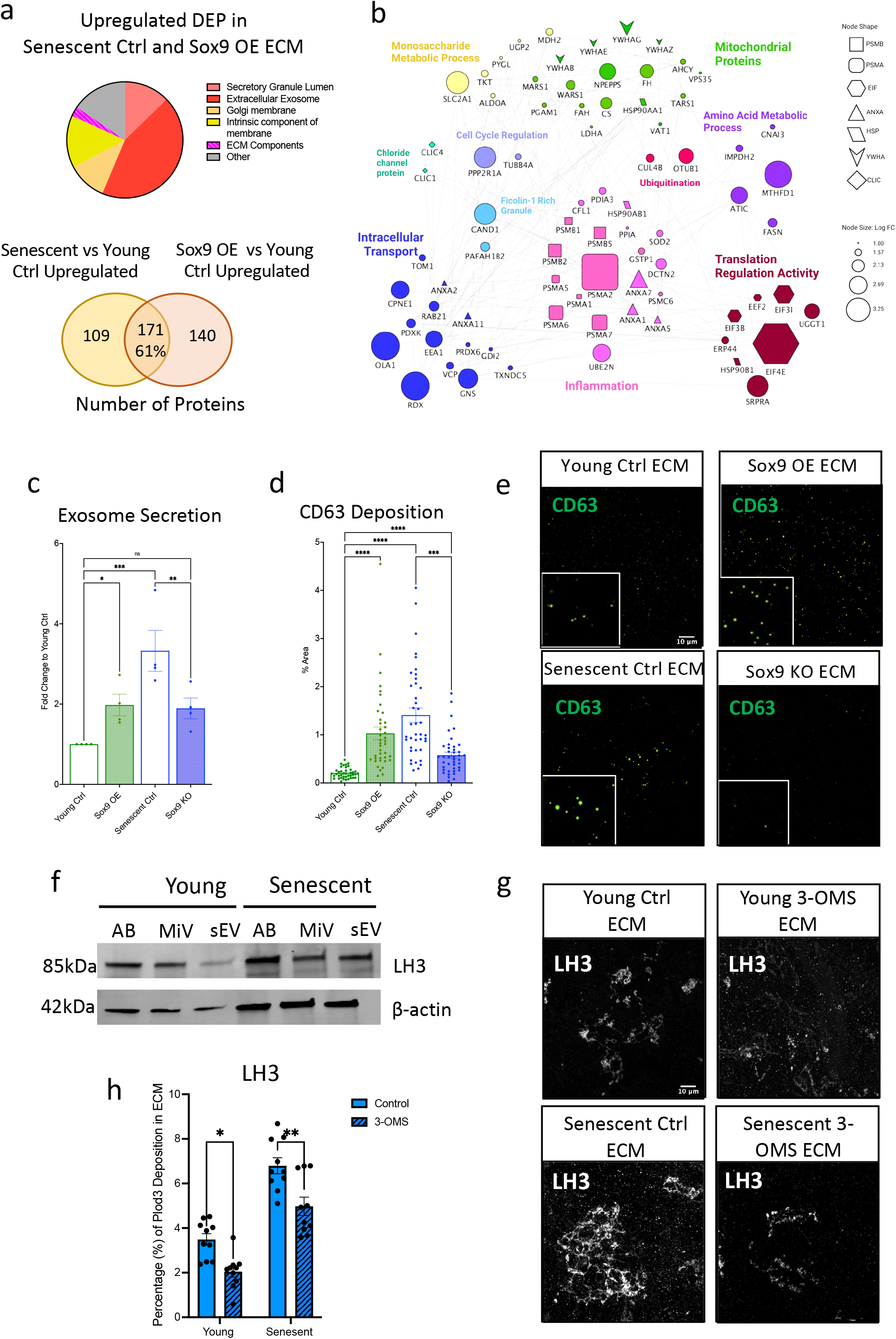
Sox9 regulates LH3 deposition into the ECM via increased extracellular vesicle (EV) secretion. (a) Pie chart and Venn-diagram depicting categories and percentages of upregulated differentially expressed proteins (DEP) found both in the ECM synthesized from senescent VSMCs transduced with EGFP (Senescent Ctrl) and young VSMCs transduced with Sox9 overexpression adenovirus (Sox9 OE). (b) Protein-protein-interaction of upregulated DEP found in Sox9 OE ECM related to extracellular vesicles through gene ontology. (c) Quantification of exosome secretion in young VSMCs transduced with EGFP control adenovirus (Young Ctrl), Sox9 OE, Senescent Ctrl, and Sox9 knockout adenovirus (Sox9 KO), n=4. (d,e) Quantification and representative IF images of CD63 deposition in the Young Ctrl, Sox9 OE, Senescent Ctrl, and Sox9 KO ECM, n=20. One-way ANOVA with Tukey post hoc test, *p-value<0.05, ***p-value<0.005, ****p-value<0.0001. (f) Western blot of LH3 protein expression in three different types of extracellular vesicles; Apoptotic bodies (AB), micro-vesicles (miV), and small EV (sEV). (g,h) Representative IF images and quantification of LH3 deposition into young and senescent ECM either with or without sEV secretion inhibitor 3-O-methylsphingomyelin (3-OMS), n=10. Two-way ANOVA, *p-value<0.05, **p-value<0.001.

## Discussion

This study explored the role and regulation of Sox9 in vascular ageing. It reveals a crucial role for Sox9 in driving the senescent ECM phenotype of VSMCs by regulating ECM stiffness, fibre alignment, and composition. Our findings suggest that elevated expression of Sox9 in VSMCs strongly depends on mechano-signalling cues from the extracellular environment. We demonstrate that Sox9 regulates ECM composition by driving a phenotype closely mimicking the senescent VSMC ECM characterized by downregulation of essential basement membrane proteins and upregulation of LH3. Crucially, these Sox9-driven ECM changes can feedback and regulate the phenotype of VSMCs by modulating proliferation, inflammation and senescence. This study also highlights the underappreciated role of LH3 as a driver of ECM stiffness and identifies EVs as novel regulators of ECM remodelling.

### Sox9 is a key regulator of VSMC phenotype during ageing

Our results have provided confirmation of previous studies that highlight the role of Sox9 in regulating VSMC contractility (22,23). We observed a negative correlation between Sox9 and α-SMA in *in vivo* human aortic tissue samples, as well as *in vitro* via Sox9 overexpression. However, our results contrast to previous studies examining the interplay between Sox9 and vascular calcification (23). We found no correlation between medial calcification and Sox9 expression *in vivo*, and we verified this *in vitro* by showing no changes in calcification propensity between control and Sox9 overexpressing VSMCs. We propose these differences may be associated with the origins of cells used in the studies. Previous studies finding an association between Sox9 and calcification were conducted in mouse models, which are genetically susceptible to spontaneous cartilaginous metaplasia that is prevalent in mice but rare in humans (36). Indeed, we found in human VSMCs that Sox9 did not increase the deposition of the chondrocyte markers Collagen 2 and 11 that have been co-localised with Sox9 in mouse aortic tissues (23). Our results are indicative of the probable interplay between Sox9 and Runx2 within the vasculature. Runx2 is a critical regulator of osteoblast differentiation (37) and a driver of vascular calcification (17,20). It also has a tight interplay with Sox9, with both TF’s exerting decreased transcriptional activation of one another (38). Thus, even though calcification and osteo/chondrogenic VSMC differentiation are tightly interconnected, the VSMCs within the microenvironments where calcification occurs potentially have a dominant Runx2 expressing phenotype, as opposed to Sox9. These findings highlight the complexity of the interplay between VSMC de-differentiation and calcification, and the need for further investigation into the regulatory mechanisms underlying these processes.

Instead, in our study, we observed a correlation between Sox9 expression and p16, a marker for cellular senescence. *In vitro* studies showed there was not a direct relationship between Sox9 levels and senescence however we found that Sox9 expression was mechanosensitive, most strongly in senescent VSMCs, with increased expression and nuclear translocation observed on stiff surfaces. This observation is in line with previous studies that have shown Sox9 is mechanosensitive in chondrocytes (30). Our findings suggest that aortic stiffening, driven in part by the stiffer modified ECM deposited by senescent VSMCs, is conducive to increased expression of Sox9 which in turn, leads to further elaboration of a senescent ECM. This may account for our observation of micro-environments in matrix-rich regions of the aorta showing localised p16 and Sox9 expression particularly, as outlined below, these ECM environments *in vitro* were shown to reduce proliferation and promote senescence.

### Sox9 induced ECM changes influence VSMC ageing and inflammation

We demonstrated the significant role of Sox9 in regulating ECM fibre alignment, ECM stiffness, and the phenotype of VSMCs. Our findings show that decreased cellular contractility, as a result of Sox9 expression, contributed to ECM fibre alignment, and we provide novel evidence that the Sox9-driven ECM compositional changes can feedback and regulate the phenotype of VSMCs. While previous studies have examined the role of ECM in cell signalling, stem cell differentiation (39,40), adhesion, and migration (41,42), the role of ECM in regenerating cells from a pathological state has remained understudied. Replating senescent cells on an ECM synthesized from Sox9-depleted VSMCs enabled, to some extent, the rejuvenation of cells with reduction in senescence markers and re-entry back into the cell cycle. Conversely VSMCs plated on ECM from Sox9-expressing cells acquired features of cellular senescence including cell cycle arrest and DNA damage. We also showed the vital role of ECM alignment in VSMC orientation. Cells plated on fibrous ECMs, such as those synthesized from senescent and young Sox9-overexpressing VSMCs, were unable to form proper alignments with neighbouring cells and fibres. Correct VSMC alignment is crucial in the vasculature, as it affects the mechanical properties of the tissue, influencing its ability to withstand and respond appropriately to mechanical stresses and strains (43). Our findings underscore the impact of Sox9-modulated ECM on the phenotype of VSMCs and further work is now required to understand how the complex compositional and structural changes contribute to VSMC ageing and rejuvenation.

Our investigation also explored the compositional changes driven by Sox9 that increase ECM stiffness. Previous studies (25,26) identified Sox9 as a potent activator of numerous ECM proteins in chondrocytes, leading us to anticipate an increase in collagen deposition. However, we found that increased Sox9 expression in VSMCs resulted in the downregulation of a wide range of collagens and glycoproteins. This observation highlights previous research showing that Sox9 can serve both as an activator and a repressor (44), as well as a regulator of both ubiquitous and chondrocyte-specific genes (45). Our findings suggest that the transcriptional targets of Sox9 vary significantly depending on the cell type in which it is expressed. In VSMCs, Sox9 acts as a repressor for the transcription of various basement membrane proteins, whose decreased expression can negatively impact both the phenotype of VSMCs (46) and the tensile strength of the vessel wall (47). Indeed, we validated both increased LH3 expression and decreased collagen IV. LH3 is essential for normal biosynthesis and secretion of type IV collagens (48) into the basement membrane and both these proteins have been implicated in vascular diseases, including Ehlers-Danlos Syndrome, which are associated with aneurysm and vessel rupture (OMIM 612394) (49). Our study suggests that loss of basement membrane integrity may also be a key feature of vascular ageing, increasing the susceptibility of the vessel to structural failure.

### LH3 and EVs as key modulators of ECM stiffening

LH3 is a novel gene regulated by Sox9 in VSMCs that plays an important role in collagen cross-linking and glycation both intra-(50) and extracellularly (51). Our results demonstrated that Sox9 and cellular senescence increased LH3 deposition in the ECM and this was confirmed *in vivo* where LH3 correlated with Sox9 and p16 expression in aged aortic samples. *In vitro* we showed that Sox9-regulated ECM stiffness was primarily driven by LH3, as its depletion resulted in a reduction of ECM stiffness, similar to that observed with the depletion of Sox9. Previous research has shown LH3 is trafficked into the extracellular space via Collagen IV carriers (35) and our analysis of EV secretion from young and senescent VSMCs revealed that LH3 is loaded onto all EV types. These findings suggest that the increased deposition of LH3 in the ECM is due to both Sox9-mediated intracellular expression and increased secretion of EVs. Importantly these carriers are able to signal remotely from the cell, suggesting that a small number of VSMCs releasing these particles may have profound effects on ECM stiffening in the local environment. This highlights LH3 and EV secretion as potential therapeutic targets for further investigation in cardiovascular pathologies.

## Experimental Procedures

### Immunohistochemistry

All human materials were handled in compliance with the Human Tissue Act (2004, UK) and with ethical approval from the research ethics committee (REC reference: 13/LO/1950). Human aortic tissue samples used are shown on Supplementary Table 1. Immunohistochemistry was performed on 7um thick paraffin-embedded human aortic tissue sections. These sections were deparaffinized, rehydrated prior to heat-mediated antigen retrieval in sodium citrate buffer. Endogenous peroxidase activity was quenched by 3% H2O2, in methanol and non-specific binding blocked with 10% serum from the source species for the secondary antibody followed by a primary antibody incubation overnight at 4 °C (Sox9, abcam, ab185966,; p16, (abcam, ab241543), α-SMA (Sigma-Aldrich’ A5691), LH3 (Proteintech, 1:100). Biotinylated secondary antibody was added, followed by avidin-biotin complex (ABC) reaction (Vector Labs kits PK-6102 & PK-6010). Staining was developed with 3, 3’-diaminobenzidine (DAB) peroxidase substrate kit (Vector Laboratories; SK-4100) and these sections were counterstained with Hematoxylin. The slides were mounted with DPX followed by dehydration and images were taken using a Leica ICC50 W Microscope, a minimum of eight images were taken of the tissue sample per aortic region (intima, media, adventitia). α-SMA and CD68, a marker of macrophage infiltration, were used to delineate the cell types present in the samples. Percentage of positively stained nuclei was quantified using ImageJ software, with brown nuclei regarded as positive and blue/purple as negative. Percentage of positive extracellular staining was quantified via threshold detection method on ImageJ.

### Cell culture

VSMCs were isolated from explants of human aortic tissues. Three different donors were used for this study; 35-year old female (04.35F.11A), 38-year old female (03.38F.11A), and 33-year old female (05.33F.5A). Cells were cultured at 37 °C and 5% CO_2_ in M199 media (Gibco), supplemented with 20% fetal bovine serum and 1% Penicillin-Streptomycin-Glutamine (PSG) (100x). Replicative senescence was induced by passaging the cells until growth arrest, typically occurring between passage 20-25. All experiments were performed using these 3 biological replicates unless stated otherwise. Early passage VSMCs (termed “young VSMCs) were collected at passage 6-13. VSMCs would reach replicative senescence in passages 25-30.

### Extracellular matrix synthesis

Decellularized ECM was synthesized as previously described (52). Plates or cover slides were coated with 0.2% filtered gelatin solution at 37°C for 1 hour, cross-linked with 1% glutaraldehyde for 30 minutes at room temperature, and quenched with 1M ethanolamine for another 30 minutes at room temperature. Cells were seeded at high confluency and cultured in 20% FBS, 1% PSG M199 media supplemented with 50ug/mL L-ascorbate for 9-12 days. The ECM was decellularized by incubating the cells for 5 minutes with the Extraction Buffer (0.1% Triton, 20mM NH4OH, diluted in PBS), then washed three times with PBS to remove cell debris.

### Adenoviral Constructs and Transfections

Sox9 was overexpressed in young VSMC via a recombinant Enhanced Green Fluorescent Protein (EGFP)-CMV-Sox9 (VectorBuilder) and knocked out in senescent VSMC via a recombinant EGFP-SOX9 shRNA (short hairpin RNA against Sox9) adenovirus (Vectorbuilder). Control young and senescent VSMCs were treated with either EGFP (VectorBuilder) or Scramble shRNA control adenovirus (VectorBuilder). Cells were transduced in serum free M199 media containing the adenovirus for 40 minutes, and then diluted in M199 media with 20% FBS and 0.1% PSG for three days. Cells synthesising matrices were re-transduced every 5 days with L-ascorbate supplementation for a total of 9 days. 100% transduction efficiency was assessed via EGFP fluorescence imaging.

### Calcification Assays

VSMCs were treated with 5% FBS supplemented M199 media containing 2.7mmol/L Calcium and 2.5mmol/L Phosphate for 8 days, when signs of calcification became apparent. Control medium was 5% FBS supplemented M199 media without added calcium and phosphate. Calcification assays were conducted as previously described (20).

### RNA Extraction, Reverse Transcription, and qPCR

RNA from cultured VSMCs was isolated using RNA STAT-60 phenol-chloroform. cDNA was synthesized using Mu-MLV reverse transcriptase with dNTPS, random and oligo primers, and RNAse inhibitor. qPCR (quantitative PCR) was performed using qPCRBIO SyGreen Mix and run in a StepOnePlus Real Time PCR system. PCRs were performed in 20uL reaction volumes, with the thermocycler program running 95 °C for 10 minutes followed by 40 cycles at 95 °C for 5 seconds and 60 °C for 30 seconds. The following primers used; 18S (QuantiTec Primer Hs-RRN18s), *Sox9* (Sox9, 5’-AGCTCTGGAGACTTCTGAACGAGA -3’, and 5’ – ACTTGTAATCCGGGTGGTCCTTCT -3’), *PLOD3* (PLOD3, 5’-TCTACACACGGCTCTACC-3’ and 5’-ACCACTTCATCTAAAGCCC-3’), *COL4A1* (COL4A1, 5’-GGCAGCCAGGACCTAAA-3’, and 5’-CCCATTCCACCAACAGAT-3’), *COL4A2* (COL4A2, 5’-ACAAGTCCTACTGGCTCTCTA-3’, and 5’-GGCCTCACACACAGAAC-3’, and *NID2 (*NID2, 5’-TTCCTTCTCCTGCCGTTGTC-3’, and 5’-TGCTGTTGTTCACAGGGTGT-3’). Expression levels of target genes were normalised to 18S expression values in young VSMC using the 2-ΔΔCt method. Ct values were normalized to those of young or senescent cells plated on plastic in hydrogel and ECM cell re-plating experiments.

### Protein Extraction and Western blotting

ECM and cell lysate was homogenized in RIPA buffer with protease inhibitor cocktail, sonicated on ice, and centrifuged for 10 minutes at 7,500 at 4°C. 50% glycerol, 0.1% SDS, 30% 1M Tris Buffer pH=6.9, Bromophenol blue 6 mM, and 10% 10mM dithiothreitol (DTT) was added to the samples prior to boiling them for 10 minutes at 95°C. Proteins were separated via SDS-PAGE, transferred to Polyvinylidine Fluoride (PVDF) membrane, and blotted with 5% BSA in PBS-Tween (0.1%) (PBS-T) for one hour at room temperature. Membranes were incubated overnight at 4°C with the primary antibodies: Sox9 (Abcam, ab185966, 1:1000), Α-SMA (Abcam, ab7818, 1:5000), LH3 (Proteintech, 11027-1AP, 1:1000). IRDye 800CW conjugated secondary antibodies (LI-COR Biosciences) were incubated for one hour at room temperature and visualized using an images (Odyssey; LI-COR Biosciences). Protein bands were quantified through ImageJ, normalized to either GAPDH or beta-actin.

### Hydrogel Setup

Hydrogel recipes measuring 10kPa and 250kPa were made up from 10.75% and 15% acrylamide and 9% and 1.2% bisacrylamide, respectively and stiffness was confirmed using AFM. Glass coverslips were coated with 3-aminopropyl trimethoxysilane (97%) for 2 minutes, washed thoroughly, and fixed for 30 minutes at room temperature with 1% glutaraldehyde with ddH_2_O. Coverslips were dried face up overnight. Per each volume of acrylamide solution, 1/100^th^ of the volume of APS (10%) and 1/1000^th^ of the volume of TEMED was added. The solution was mixed thoroughly and aliquoted out onto glass slides. The fixed coverslips were placed onto the hydrogel solution facedown. The hydrogel solution was allowed to solidify and adhere onto the glass coverslips. The glass coverslips were then inverted, hydrogel slide up, and subjected to ultraviolet light (UV) for 5 minutes with a 2% of sulfa SANPAH solution (Thermo scientific, 22589). The hydrogels were washed several times under sterile conditions and incubated with 3.3% of Rat Tail Collagen 1 (Gibco A10483) solution at 4°C for 2 hours. The collagen solution was then removed and cells were plated onto the hydrogels as normal.

### Immunofluorescence

VSMCs and ECM were fixed in 4% paraformaldehyde in PBS, and permeabilized with 0.5% PF-40. Permeabilization step was omitted in case of ECM staining. Cells were then blocked at room temperature with 3% BSA in PBS, and incubated with the primary antibodies; Fibronectin (ab2413, 1;500), Sox9 (ab185966, 1:100), CD63, (Santa Cruz, s-15,363, 1:200), LH3 (Proteintech, 11027-1-AP, 1:250), Vinculin (Sigma Aldrich,V9264, 1:500), gamma-H2AX (Cell Signaling, 2577,1:200) diluted in the blocking solution overnight at 4°C. Cells were then washed and incubated in the dark with the fluorophore conjugated secondary antibody (Donkey anti-mouse/ rabbit, A10036/ A10040, 1:400). Cells were incubated with Phalloidin (Sigma Aldrich, 1:400), diluted in PBS, post-secondary antibody staining. The nuclear staining was obtained with DAPI (Sigma Aldrich, D9542) at room temperature. Coverslips were mounted onto slides using Mowiol media and visualized using Z-stacks of ECM and VSMCs was acquired using the Nikon Eclipse Ti Inverted Spinning Disk Confocal System. The images and fluorescence intensity were analysed by calculating the corrected total cell fluorescence (CTCF) on FIJI.

ECM fiber and cellular alignment analysis was performed using the Alignment by Fourier Transform (AFT) tool as previously described (53). This algorithm divides inputted images into a grid of overlapping windows of a specified size, employing Fast Fourier Transforms (FFT) on each window to construct a vector field of the entire image which is analysed in frequency space. For each individual window vector, the angle with respect to a surrounding grid of vectors is then calculated, with the size of this grid specified by a neighbourhood radius. Based upon the relative angles of these vectors, an alignment score or ‘order parameter’ is calculated between -1 and 1 for each vector, where -1 represents completely orthogonal alignment, 0 represents random alignment, and 1 represents completely parallel alignment. ECM and VSMCs were visualised by staining for fibronectin and vinculin respectively. To analyse fibronectin images with sufficient sensitivity to detect the local structural variation in the ECM meshwork, while maintaining comparison of fibre alignment over the greatest possible area, a small window of 50 px and large neighbourhood radius of 7 were used. Conversely, in VSMC images where granularity is comparatively low, a large window of 250 px and small neighbourhood radius of 1 were utilised, in order to capture the directionality only of whole cells and not finer structures or edges. Window overlap was 50% in all analyses.

### Edu Cell Proliferation

To measure cell proliferation, VSMCs were depleted from FBS for 24h to restart the cell cycle. Following depletion, cells were incubated with 10uM Click-iT Edu (ThermoFisher, C10337) for 1 hour in 20% FBS, 1% PSG supplemented M199 media. Edu visualization was achieved according to manufacturer protocol prior to any further immunofluorescence staining.

### CD63 Beads Assay

Quantification of CD63 positive sEV secretion was performed by the CD63 bead assay and flow cytometry analysis as previously described (54). 1x10^8^ 4μm aldehyde-sulphate beads (Invitrogen, A37304) were added to filter sterilised 150μl MES buffer (Sigma, 76039). Beads were spun for 10 minutes at 3000g and washed twice more with MES buffer. Beads were incubated with 35μg of CD63 antibodies (BD Biosciences, 556019) overnight at room temperature with shaking. The beads were washed thrice with 4% filter sterilised BSA-PBS and resuspended in storage buffer (0.1% Glycine, 0.1% NaN_3_ in PBS). Young and senescent VSMC were plated at a concentration of 15,000 cells per well. The following day they were washed thoroughly to remove residual exosomes and transduced with the control EGFP, Sox9 overexpression and, knockout adenovirus in 2.5% exosome-free FBS M199 media for three days. The media was then collected, spun for 5 minutes at 2500g and transferred to a new 24-well plate. The bead stock solution was washed once with 2% BSA-PBS, with 1μL of bead suspension per each media sample. Beads were incubated in the media overnight at 4°C with gentle rocking. Cells were trypsinized and incubated with Solution13

(ChemoMetec) for a cell viability assay using the NC3000 cell nucleocounter (ChemoMetec). The beads were washed twice with 2% BSA-PBS and incubated with phycoerythrin (PE)-conjugated light sensitive anti-CD81 antibody for 1 hour at room temperature. The beads were quantified using FACs analysis with a BD Accuri C6 flow cytometer. Data analysis was conducted using FlowJo software, with the number of sEVs normalised to the cell number that was calculated during the cell viability assay.

### Proteomics and bioinformatic analysis

Decellularized ECM protein samples were de-glycosylated as previously described (55) Samples were precipitated overnight in acetone at -20°C and the protein pellet airdried in a speed vacuum. They were then incubated with chondroitinase, heparinase, keratanase, and 3 de-branching enzymes +1 O-deglycosidase diluted in the de-glycosylation buffer (Tris 200mM, NaAc 200mM, EDTA 100mM) at 25°C for two hours and then at 37°C overnight with shaking. Samples were air dried once again and incubated with PNGaseF diluted in H_2_0^18^ for 48 hours at 37°C with shaking. Sample purification and trypsin digestion was then undertaken for peptide extraction. Samples were injected to be analyzed by LC-MS/MS in triplicate. Raw mass spectrometry data was processed into peak list files using Proteome Discoverer (ThermoScientific, v2.5) and processed using Sequest (Eng et al; PMID 24226387) search algorithms against the Uniprot Human Taxonomy database with a 1% FDR stringency. The overall intensity value for each peptide was compared across the sample set to further calculate changes in protein abundance. Data has been uploaded to PRIDE. Accession numbers ( ).

Differentially expressed proteins were identified using DEP and Limma packages in R (Fold change set to 1, minimum P-value 0.05). David Gene Ontology (GO) was used to identify the categories that the differentially expressed proteins belonged to. Proteins belonging to the classification of “extracellular space”, “extracellular region”, “basement membrane”, and “extracellular matrix” were extracted, pooled together, and further sub-classified in Cytoscape app 3.8.0 with GeneMANIA app. The same process was repeated for all proteins initially classified under “extracellular exosome”. Protein-protein interaction maps were built in Cytoscape, with proteins exhibiting minimal interactions with the network being discarded.

### Atomic Force Microscopy

Atomic Force Microscopy (AFM) data acquisition was done on JPK Nanowizard 3. For AFM imaging, a tetrahedral shaped cantilever with a silicone tip with a spring constant of 0.081 M/m, tip height of 4-5μm (HYDRA-6v-200WG, Applied NanoStructures, CA) was used. The matrix stiffness was recorded via Contact Mode, measuring the force spectroscopy, in liquid. The set point was 5nN with a Z-length of 14 microns and extend speed of 10μm/second. The AFM data was processed via the JPK Nanowizard 3 data processing software.

### Vesicle Isolation

EVs were isolated and purified by ultracentrifugation (UC) as previously described (54). VSMCs were washed twice with Earl’s balanced salt solution (EBSS) and the media changed to Dulbecco’s modified eagle medium (DMEM) supplemented with 0.1% BSA and 1% PSG. After 24h the media was collected and centrifuged at 1000g to pellet apoptotic bodies, at 10,000g to pellet micro-vesicles, and 100,000g by UC to pellet small EVs. Pellets were washed and resuspended in PBS.

### Statistical Analysis

All results are from three different VSMC donor isolates. All results are presented as +/- Standard error of mean (SEM). Statistical analysis was performed in GraphPad software. Normality tests were conducted for all datasets via the Shapiro-Wilk test prior to comparison analysis. Multiple groups were compared via the 1-way ANOVA with Tukey test, whilst two independent groups were compared via the Student *t* test. Simple linear correlation was conducted to determine the p value and R squared fit between two different proteins in histological samples. *p-value< 0.05, **p-value<0.01, ***p-value<0.001, ****p-value<0.0001.

## Supplementary Materials

S Table 1: Table describing the age, gender, American Heart Association (AHA) classification of atherosclerotic plaques, and level of calcification within the Medial aortic area. Lack of calcification is denoted by -, whist severity of calcification is graded from + to + + + +.

S Fig1: Sox 9 does not associate with vascular calcification. (a) Representative images of matched patient tissue samples stained for Von Kossa (Calcification), Sox9, and p16. Red arrows indicate prominent positive cellular staining. Correlation between calcium deposition (Medial Calcification %), age, cellular senescence (P16), and Sox9 expression. Linear correlation between (b) Medial Calcification and Patient Age, (c) Medial Calcification and positive p16 (%) cells, and (d) Medial calcification and positive Sox9 (%) cells (e) Sox9 and patient age. Statistical significance determined via Simple Linear Regression. (f) Alizarin red staining showing Sox9 did not increase calcification of VSMCs *in vitro*. (g) RT-qPCR showing Sox9 expression did not change osteogenic gene expression in calcifying VSMCs.

S Fig2: Sox9 regulates Vascular Smooth Muscle Cell Phenotype. (a) Gene expression, quantified via RT-qPCR of TAGLN, in young VSMCs transfected with control EGFP adenovirus (Young Ctrl) and Sox9 overexpressing adenovirus (Sox9 OE), n=4. (b,c) Quantification of protein expression and Western blot of *ACTA2*/α-SMA from young Ctrl and Sox9 OE cell lysate samples, n=4 (d) Western blot of Sox9 expression in young Ctrl, Sox9 OE, senescent VSMCs transduced with shEGFP control adenovirus (Senescent Ctrl), and Sox9 knockout adenovirus (Sox9 KO). (e) Gene expression, quantified by RT-qPCR, of Sox9 in senescent VSMCs transduced with control shEGFP adenovirus (Senescent Ctrl) and Sox9 knockout (Sox9 KO) adenovirus, n=4, Unpaired Student’s T-test, *p-value<0.05. (f) Gene expression, quantified by RT-qPCR, of p16 in Young Ctrl, Sox9 OE, Senescent Ctrl, and Sox9 KO, n=4. Gene expression, quantified by RT-qPCR, of Sox5 in (g) young and (h) senescent VSMCs plated on matrices of different stiffness and plastic coated with collagen (P+C). Gene expression, quantified by RT-qPCR, of Sox8 of (i) young and (j) senescent VSMCs plated on hydrogels of 10kpa and 250kpa, and P+C, n=4. Genes normalized to young and senescent VSMCs plated on plastic (indicated by the dotted line).One-way ANOVA with Tukey post hoc test, *p-value<0.05.

S. Figure 3: Gene expression of *p16* in (a) young and (b) senescent VSMCs when plated on hydrogels of stiffness 10kPa and 250kpa, and plastic coated with collagen (P+C). Gene expression, of *p21* in (c) young and (d) senescent VSMCs when plated on hydrogels of stiffness 10kPa and 250kpa, and P+C. Gene expression of *IL6* in (e) young and (f) senescent VSMCs when plated on hydrogels of stiffness 10kPa and 250kpa, and P+C. Gene expression of *ACTA2* in (g) young and (h) senescent VSMCs when plated on hydrogels of stiffness 10kPa and 250kpa, and P+C. Gene expression of *TAGLN* in (i) young and (j) senescent VSMCs when plated on hydrogels of stiffness 10kPa and 250kpa, and P+C. N=7, One-way ANOVA with Tukey post hoc test, *p-value<0.05. Gene expression of (k) *ACTA2* and (l) *TAGLN* from young and senescent VSMCs plated on ECM synthesized from young VSMCS treated with EGFP control adenovirus (Young Ctrl), Sox9 overexpressing adenovirus (Sox9 OE), senescent VSMCs treated with shEGFP control adenovirus (Senescent Ctrl), and Sox9 knockout adenovirus (Sox9 KO). N=4, Unpaired Student’s t-test. All gene expressions were quantified via RT-qPCR.

S Figure 4: (a) Principal component Analysis (PCA) plot of proteomics samples. Circle indicates clustered cell populations. (b) GO analysis showing pathways regulated by differentially expressed genes between young and senescent VSMCs. Gene expression, quantified via RT-qPCR of (c) Collagen (Col)4a1, (d), Col4a2, and (e) nidogen2 (Nid2) from young VSMCs transduced with EGFP control adenovirus (Young Ctrl), Sox9 overexpression adenovirus (Sox9 OE), and senescent cells transduced with shEGFP control adenovirus (Senescent Ctrl). N=4, One-way ANOVA with Tukey post hoc test, **p-value<0.01, ***p-value<0.005, p****<0.00001. (f) Immunofluorescence of Collagen 4 (Col IV) in ECM synthesized from Young Ctrl, Sox9 OE, Senescent Ctrl, and senescent VSMCs transduced with Sox9 knockout adenovirus (Sox9 KO).

S Figure 5: (a) Representative immunofluorescence images of LH3 in young VSMCs transduced with EGFP control adenovirus (Young Ctrl), Sox9 overexpression (Sox9 OE), and senescent VSMCs transduced with shEGFP control adenovirus (Senescent Ctrl), and Sox9 knockout (Sox9 KO). LH3 is in grey and red, EGFP in green, and nuclear staining (DAPI) in blue. (b) Western blot depicting LH3 knockout (siPlod3) with Sox9 overexpression (Sox9 OE) and Sox9 knockout (KO) from protein lysate in young and senescent Vascular Smooth Muscle Cells.

S. Figure 6: Protein quantification of LH3 from three types of extracellular vesicles in young and senescent Vascular Smooth muscle cells.

## Author Contributions

MF, SC and CMS contributed to conception; MF, MW, SC and CMS to experimental design; MF, SA and GW to acquisition of data; MF and SA to analysis and interpretation of data. MF and CMS wrote and revised the manuscript and all authors provided final approval of the submitted version.

## Acknowledgements

This study was supported by the British Heart Foundation programme grant RG/F/21/110064 to CS and KCL BHF Centre of Research Excellence Interdisciplinary PhD studentship to CS and SC. The microscopy was performed at the Wohl Cellular Imaging Centre at King’s College London. The proteomics was performed at the Denmark Hill Proteomics Facility. Graphical abstract and schematic created with BioRender.com.

## Conflict of Interest

The authors declare that they have no conflicts of interest.

## Notes

### Competing Interest Statement

The authors have declared no competing interest.

### Summary of Updates

This version of the manuscript has been revised to include new data analysis and to update the discussion and citations.

## References

1. Lanzer P, Boehm M, Sorribas V, Thiriet M, Janzen J, Zeller T, et al. Medial vascular calcification revisited: review and perspectives. Eur Heart J. 2014 Jun 14;35(23):1515–25.

2. Speer MY, Yang HY, Brabb T, Leaf E, Look A, Lin WL, et al. Smooth Muscle Cells Give Rise to Osteochondrogenic Precursors and Chondrocytes in Calcifying Arteries. Circ Res. 2009 Mar 27;104(6):733–41.

3. Naik V, Leaf EM, Hu JH, Yang HY, Nguyen NB, Giachelli CM, et al. Sources of cells that contribute to atherosclerotic intimal calcification: an in vivo genetic fate mapping study. Cardiovasc Res. 2012 Jun 1;94(3):545–54.

4. Bobryshev Y V. Transdifferentiation of smooth muscle cells into chondrocytes in atherosclerotic arteries in situ: implications for diffuse intimal calcification. J Pathol. 2005 Apr 1;205(5):641–50.

5. Tyson KL, Reynolds JL, McNair R, Zhang Q, Weissberg PL, Shanahan CM. Osteo/Chondrocytic Transcription Factors and Their Target Genes Exhibit Distinct Patterns of Expression in Human Arterial Calcification. Arterioscler Thromb Vasc Biol. 2003 Mar 1;23(3):489–94.

6. Durham AL, Speer MY, Scatena M, Giachelli CM, Shanahan CM. Role of smooth muscle cells in vascular calcification: implications in atherosclerosis and arterial stiffness. Cardiovasc Res. 2018 Mar 3;114(4):590.

7. Reynolds JL, Joannides AJ, Skepper JN, Mcnair R, Schurgers LJ, Proudfoot D, et al. Human vascular smooth muscle cells undergo vesicle-mediated calcification in response to changes in extracellular calcium and phosphate concentrations: A potential mechanism for accelerated vascular calcification in ESRD. J Am Soc Nephrol. 2004 Nov;15(11):2857–67.

8. Nakano-Kurimoto R, Ikeda K, Uraoka M, Nakagawa Y, Yutaka K, Koide M, et al. Replicative senescence of vascular smooth muscle cells enhances the calcification through initiating the osteoblastic transition. Am J Physiol Hear Circ Physiol. 2009;297:1673–84.

9. Liu Y, Drozdov I, Shroff R, Beltran L, Shanahan C. Prelamin A accelerates vascular calcification via activation of the DNA damage response and senescence-associated secretory phenotype in vascular smooth muscle cells. Circ Res. 2013;112(10).

10. Sanchis P, Ho CY, Liu Y, Beltran LE, Ahmad S, Jacob AP, et al. Arterial “inflammaging” drives vascular calcification in children on dialysis. Kidney Int. 2019 Apr 4;95(4):958.

11. Wallis R, Mizen H, Bishop CL. The bright and dark side of extracellular vesicles in the senescence-associated secretory phenotype. Mech Ageing Dev. 2020 Jul 1;189:111263.

12. Coppé JP, Desprez PY, Krtolica A, Campisi J. The Senescence-Associated Secretory Phenotype: The Dark Side of Tumor Suppression. Annu Rev Pathol. 2010 Feb 2;5:99.

13. Alique M, Bodega G, Corchete E, García-Menéndez E, de Sequera P, Luque R, et al. Microvesicles from indoxyl sulfate-treated endothelial cells induce vascular calcification in vitro. Comput Struct Biotechnol J. 2020;18:953–66.

14. Duca L, Blaise S, Romier B, Laffargue M, Gayral S, El Btaouri H, et al. Matrix ageing and vascular impacts: focus on elastin fragmentation. Cardiovasc Res. 2016 Jun 1;110(3):298–308.

15. Mammoto A, Matus K, Mammoto T. Extracellular Matrix in Aging Aorta. Front Cell Dev Biol. 2022 Feb 21;10:367.

16. Chen H, Tan XN, Hu S, Liu RQ, Peng LH, Li YM, et al. Molecular Mechanisms of Chondrocyte Proliferation and Differentiation. Front Cell Dev Biol. 2021 May 28;9:664168.

17. Lin M-E, Chen TM, Wallingford MC, Nguyen NB, Yamada S, Sawangmake C, et al. Runx2 deletion in smooth muscle cells inhibits vascular osteochondrogenesis and calcification but not atherosclerotic lesion formation. Cardiovasc Res. 2016 Aug 5;112:606–16.

18. Sun Y, Byon CH, Yuan K, Chen J, Mao X, Heath JM, et al. Smooth muscle cell-specific runx2 deficiency inhibits vascular calcification. Circ Res. 2012 Aug 17;111(5):543–52.

19. Duer M, Cobb AM, Shanahan CM. DNA Damage Response: A Molecular Lynchpin in the Pathobiology of Arteriosclerotic Calcification. Arterioscler Thromb Vasc Biol. 2020 Jul 1;40:E193–202.

20. Cobb AM, Yusoff S, Hayward R, Ahmad S, Sun M, Verhulst A, et al. Runx2 (Runt-Related Transcription Factor 2) Links the DNA Damage Response to Osteogenic Reprogramming and Apoptosis of Vascular Smooth Muscle Cells. Arterioscler Thromb Vasc Biol. 2021 Apr 1;41(4):1339–57.

21. Briot A, Jaroszewicz A, Warren CM, Lu J, Touma M, Rudat C, et al. Repression of Sox9 by Jag1 Is Continuously Required to Suppress the Default Chondrogenic Fate of Vascular Smooth Muscle Cells. Dev Cell. 2014 Dec 22;31(6):707–21.

22. Xu Z, Ji G, Shen J, Wang X, Zhou J, Li L. SOX9 and myocardin counteract each other in regulating vascular smooth muscle cell differentiation. Biochem Biophys Res Commun. 2012 Jun 1;422(2):285–90.

23. Augstein A, Mierke J, Poitz DM, Strasser RH. Sox9 is increased in arterial plaque and stenosis, associated with synthetic phenotype of vascular smooth muscle cells and causes alterations in extracellular matrix and calcification. Biochim Biophys Acta - Mol Basis Dis. 2018 Aug 1;1864(8):2526–37.

24. Yu Q, Li W, Xie D, Zheng X, Huang T, Xue P, et al. PI3Kγ promotes vascular smooth muscle cell phenotypic modulation and transplant arteriosclerosis via a SOX9-dependent mechanism. EBioMedicine. 2018 Oct 1;36:39–53.

25. Lefebvre V, Huang W, Harley VR, Goodfellow PN, de Crombrugghe B. SOX9 is a potent activator of the chondrocyte-specific enhancer of the pro alpha1(II) collagen gene. Mol Cell Biol. 1997 Apr;17(4):2336–46.

26. Oh C Do, Lu Y, Liang S, Mori-Akiyama Y, Chen D, De Crombrugghe B, et al. SOX9 Regulates Multiple Genes in Chondrocytes, Including Genes Encoding ECM Proteins, ECM Modification Enzymes, Receptors, and Transporters. PLoS One. 2014 Sep 17;9(9):e107577.

27. Gridley T. Notch signaling in vascular development and physiology. Development. 2007 Aug 1;134(15):2709–18.

28. Seime T, Akbulut AC, Liljeqvist ML, Siika A, Jin H, Winski G, et al. Proteoglycan 4 modulates osteogenic smooth muscle cell differentiation during vascular remodeling and intimal calcification. Cells. 2021 Jun 1;10(6):1276.

29. Tyson KL, Reynolds JL, McNair R, Zhang Q, Weissberg PL, Shanahan CM. Osteo/chondrocytic transcription factors and their target genes exhibit distinct patterns of expression in human arterial calcification. Arterioscler Thromb Vasc Biol. 2003 Mar 1;23(3):489–94.

30. Woods A, Wang G, Beier F. RhoA/ROCK signaling regulates Sox9 expression and actin organization during chondrogenesis. J Biol Chem. 2005 Mar 25;280(12):11626– 34.

31. Porter LJ, Holt MR, Soong D, Shanahan CM, Warren DT. Prelamin A Accumulation Attenuates Rac1 Activity and Increases the Intrinsic Migrational Persistence of Aged Vascular Smooth Muscle Cells. Cells 2016, Vol 5, Page 41. 2016 Nov 15;5(4):41.

32. Kumari R, Jat P. Mechanisms of Cellular Senescence: Cell Cycle Arrest and Senescence Associated Secretory Phenotype. Front Cell Dev Biol. 2021 Mar 29;9:645593.

33. Kapustin AN, Davies JD, Reynolds JL, McNair R, Jones GT, Sidibe A, et al. Calcium regulates key components of vascular smooth muscle cell-derived matrix vesicles to enhance mineralization. Circ Res. 2011 Jun 24;109(1).

34. Mas-Bargues C, Borrás C, Alique M. The Contribution of Extracellular Vesicles From Senescent Endothelial and Vascular Smooth Muscle Cells to Vascular Calcification. Front Cardiovasc Med. 2022 Apr 15;9.

35. Banushi B, Forneris F, Straatman-Iwanowska A, Strange A, Lyne AM, Rogerson C, et al. Regulation of post-Golgi LH3 trafficking is essential for collagen homeostasis. Nat Commun. 2016 Jul 20;7.

36. Qiao JH, Fishbein MC, Demer LL, Lusis AJ. Genetic Determination of Cartilaginous Metaplasia in Mouse Aorta. Arterioscler Thromb Vasc Biol. 1995;15(12):2265–72.

37. Takarada T, Hinoi E, Nakazato R, Ochi H, Xu C, Tsuchikane A, et al. An analysis of skeletal development in osteoblast-specific and chondrocyte-specific runt-related transcription factor-2 (Runx2) knockout mice. J Bone Miner Res. 2013 Oct;28(10):2064–9.

38. Cheng A, Genever PG. SOX9 determines RUNX2 transactivity by directing intracellular degradation. J Bone Miner Res. 2010;25(12):2680–9.

39. Watt FM, Huck WTS. Role of the extracellular matrix in regulating stem cell fate. Nat Rev Mol Cell Biol 2013 148. 2013 Jul 10;14(8):467–73.

40. Hoshiba T, Chen G, Endo C, Maruyama H, Wakui M, Nemoto E, et al. Decellularized Extracellular Matrix as an In Vitro Model to Study the Comprehensive Roles of the ECM in Stem Cell Differentiation. Stem Cells Int. 2016;2016.

41. Rickel AP, Sanyour HJ, Leyda NA, Hong Z. Extracellular Matrix Proteins and Substrate Stiffness Synergistically Regulate Vascular Smooth Muscle Cell Migration and Cortical Cytoskeleton Organization. ACS Appl Bio Mater. 2020 Apr 20;3(4):2360– 9.

42. Scherzer MT, Waigel S, Donninger H, Arumugam V, Zacharias W, Clark G, et al. Fibroblast-Derived Extracellular Matrices: An Alternative Cell Culture System That Increases Metastatic Cellular Properties. PLoS One. 2015 Sep 15;10(9):e0138065.

43. Zhu M, Wang Z, Zhang J, Wang L, Yang X, Chen J, et al. Circumferentially aligned fibers guided functional neoartery regeneration in vivo. Biomaterials. 2015 Aug 1;61:85–94.

44. Lefebvre V, Dvir-Ginzberg M, Hebrew : SOX9 and the many facets of its regulation in the chondrocyte lineage HHS Public Access. Connect Tissue Res. 2017;58(1):2–14.

45. Ohba S, He X, Hojo H, McMahon AP. Distinct Transcriptional Programs Underlie Sox9 Regulation of the Mammalian Chondrocyte. Cell Rep. 2015 Jul 14;12(2):229– 43.

46. Mao C, Ma Z, Jia Y, Li W, Xie N, Zhao G, et al. Nidogen-2 Maintains the Contractile Phenotype of Vascular Smooth Muscle Cells and Prevents Neointima Formation via Bridging Jagged1-Notch3 Signaling. Circulation. 2021 Oct 12;144:1244–61.

47. Steffensen LB, Stubbe J, Lindholt JS, Beck HC, Overgaard M, Bloksgaard M, et al. Basement membrane collagen IV deficiency promotes abdominal aortic aneurysm formation. Sci Reports |. 2021;11:12903.

48. Salo AM, Cox H, Farndon P, Moss C, Grindulis H, Risteli M, et al. A Connective Tissue Disorder Caused by Mutations of the Lysyl Hydroxylase 3 Gene. Am J Hum Genet. 2008 Oct 10;83(4):495.

49. Steffensen LB, Stubbe J, Lindholt JS, Beck HC, Overgaard M, Bloksgaard M, et al. Basement membrane collagen IV deficiency promotes abdominal aortic aneurysm formation. Sci Reports 2021 111. 2021 Jun 18;11(1):1–13.

50. Myllylä R, Wang C, Heikkinen J, Juffer A, Lampela O, Risteli M, et al. Expanding the lysyl hydroxylase toolbox: New insights into the localization and activities of lysyl hydroxylase 3 (LH3). J Cell Physiol. 2007 Aug 1;212(2):323–9.

51. Salo AM, Wang C, Sipilä L, Sormunen R, Vapola M, Kervinen P, et al. Lysyl hydroxylase 3 (LH3) modifies proteins in the extracellular space, a novel mechanism for matrix remodeling. J Cell Physiol. 2006 Jun 1;207(3):644–53.

52. Beacham DA, Amatangelo MD, Cukierman E. Preparation of Extracellular Matrices Produced by Cultured and Primary Fibroblasts. Curr Protoc Cell Biol. 2006 Dec 1;33(1):10.9.1-10.9.21.

53. Marcotti S, Belo de Freitas D, Troughton LD, Kenny FN, Shaw TJ, Stramer BM, et al. A workflow for rapid unbiased quantification of fibrillar feature alignment in biological images. Front Comput Sci. 2021 Oct 14;3.

54. Whitehead M, Yusoff S, Ahmad S, Schmidt L, Mayr M, Madine J, et al. Vascular smooth muscle cell senescence accelerates medin aggregation via small extracellular vesicle secretion and extracellular matrix reorganization. Aging Cell. 2023 Feb 1;22(2):e13746.

55. Barallobre-Barreiro J, Radovits T, Fava M, Mayr U, Lin WY, Ermolaeva E, et al. Extracellular Matrix in Heart Failure: Role of ADAMTS5 in Proteoglycan Remodeling. Circulation. 2021 Dec 21;144(25):2021–34.

