## Supplemental Figures for "Sox9 accelerates vascular ageing by regulating extracellular matrix composition and stiffness"

| Age | Gender | AHA classification of atherosclerotic plaque | Calcification |
|---|---|---|---|
| 18 | M | I | - |
| 19 | M | I | - |
| 23 | M | I | + + |
| 30 | F | III | - |
| 34 | M | II | + + + + |
| 38 | F | II | + |
| 39 | M | III | +++ |
| 41 | M | II | + + + |
| 44 | M | II | - |
| 45 | M | III | - |
| 47 | M | III | + + + + |
| 49 | F | III | +++ |
| 50 | M | IV | + |
| 55 | F | II | + + + |
| 58 | M | III | + + + + |
| 63 | F | IV | + + + + |
| 66 | M | II | - |
| 70 | M | IV | ++ |
| 84 | F | III | + + + + |
| 85 | F | IV | - |
| 89 | F | IV | + + |

S Table 1: Table describing the age, gender, American Heart Association (AHA) classification of atherosclerotic plaques, and level of calcification within the Medial aortic area. Lack of calcification is denoted by - , whist severity of calcification is graded from + to + + + +.

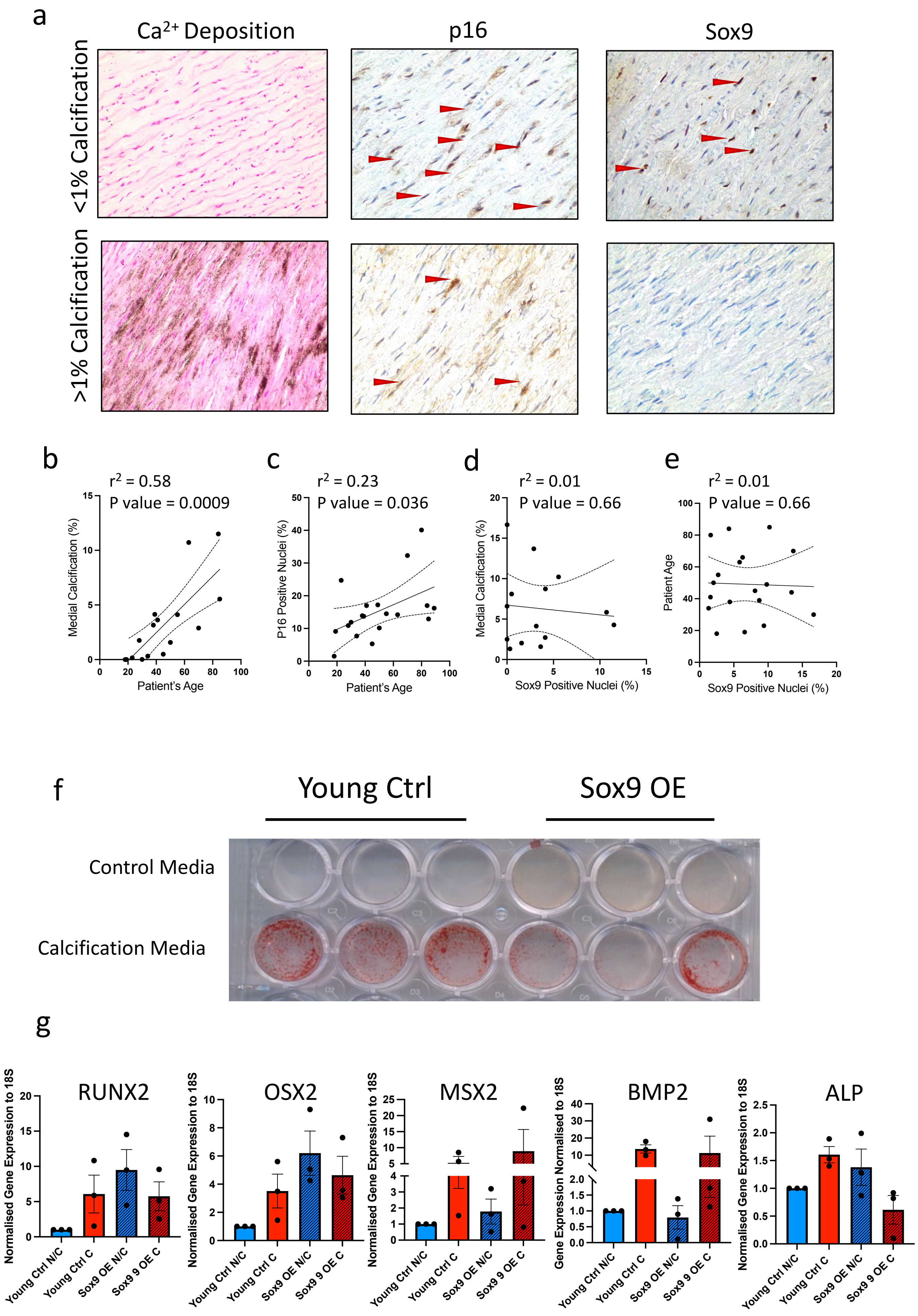


S Fig1: Sox 9 does not associate with vascular calcification. (a) Representative images of matched patient tissue samples stained for Von Kossa (Calcification), Sox9, and p16. Red arrows indicate prominent positive cellular staining. Correlation between calcium deposition (Medial Calcification %), age, cellular senescence (P16), and Sox9 expression. Linear correlation between (b) Medial Calcification and Patient Age, (c) Medial Calcification and positive p16 (%) cells, and (d) Medial calcification and positive Sox9 (%) cells (e) Sox9 and patient age. Statistical significance determined via Simple Linear Regression. (f) Alizarin red staining showing Sox9 did not increase calcification of VSMCs *in vitro*. (g) RT-qPCR showing Sox9 expression did not change osteogenic gene expression in calcifying VSMCs.

S. Figure 1

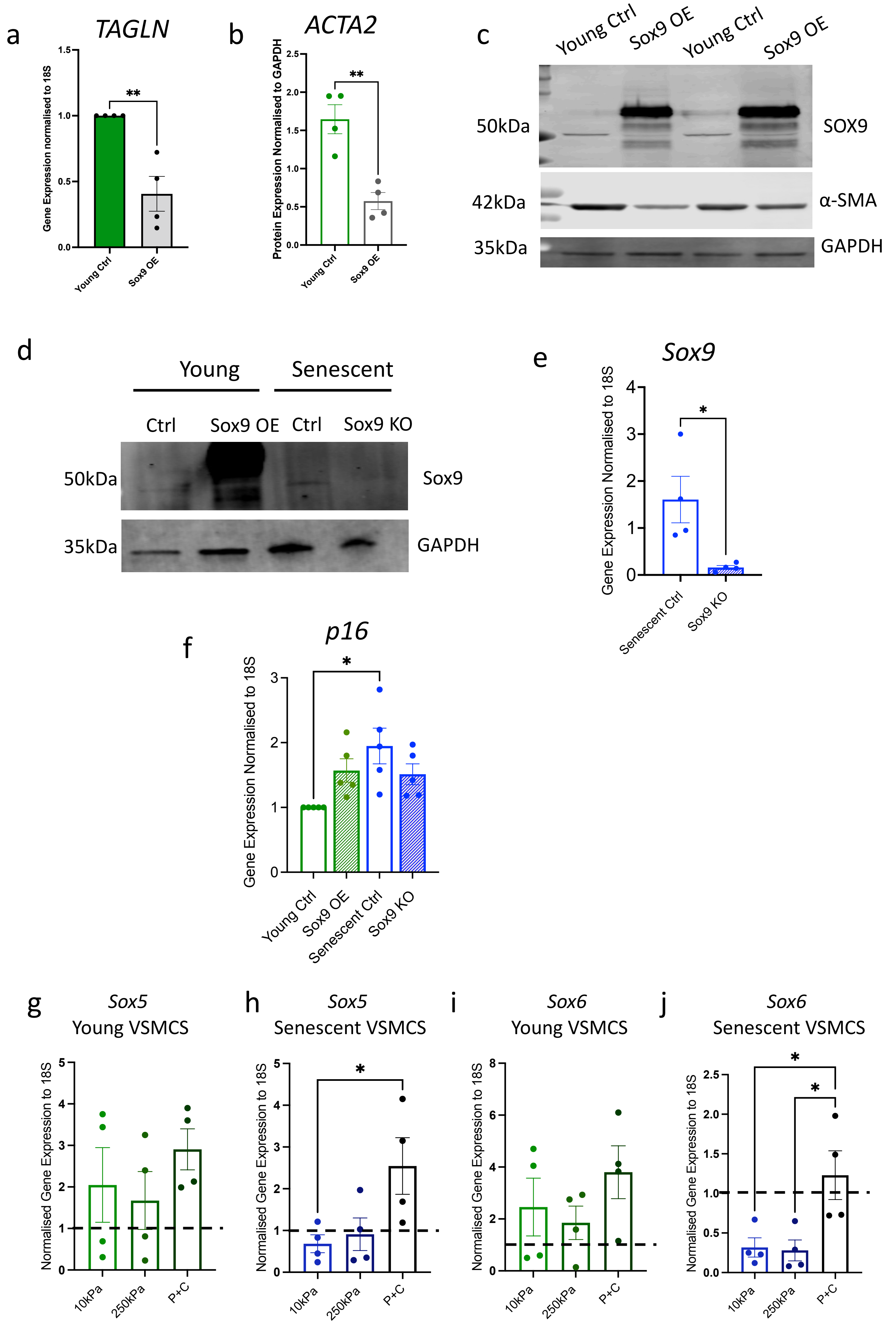


S Fig2: Sox9 regulates Vascular Smooth Muscle Cell Phenotype. (a) Gene expression, quantified via RT-qPCR of TAGLN, in young VSMCs transfected with control EGFP adenovirus (Young Ctrl) and Sox9 overexpressing adenovirus (Sox9 OE), n=4. (b,c) Quantification of protein expression and Western blot of *ACTA2*/α-SMA from young Ctrl and Sox9 OE cell lysate samples, n=4 (d) Western blot of Sox9 expression in young Ctrl, Sox9 OE, senescent VSMCs transduced with shEGFP control adenovirus (Senescent Ctrl), and Sox9 knockout adenovirus (Sox9 KO). (e) Gene expression, quantified by RT-qPCR, of Sox9 in senescent VSMCs transduced with control shEGFP adenovirus (Senescent Ctrl) and Sox9 knockout (Sox9 KO) adenovirus, n=4, Unpaired Student's T-test, *p-value<0.05. (f) Gene expression, quantified by RT-qPCR, of p16 in Young Ctrl, Sox9 OE, Senescent Ctrl, and Sox9 KO, n=4. Gene expression, quantified by RT-qPCR, of Sox5 in (g) young and (h) senescent VSMCs plated on matrices of different stiffness and plastic coated with collagen (P+C). Gene expression, quantified by RT-qPCR, of Sox8 of (i) young and (j) senescent VSMCs plated on hydrogels of 10kpa and 250kpa, and P+C, n=4. Genes normalized to young and senescent VSMCs plated on plastic (indicated by the dotted line).One-way ANOVA with Tukey post hoc test, *p-value<0.05.

S. Figure 2

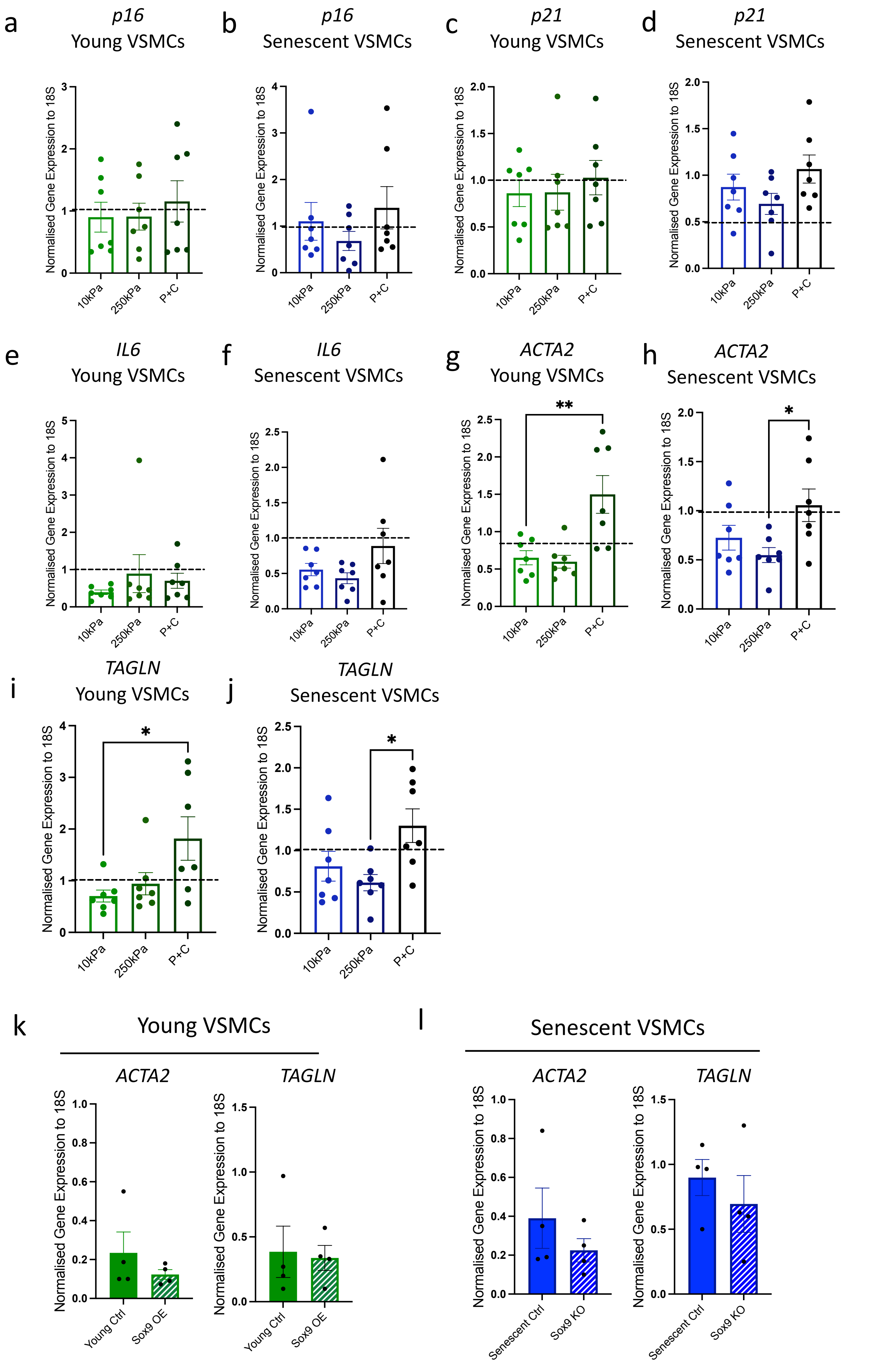


S. Figure 3: Gene expression of *p16* in (a) young and (b) senescent VSMCs when plated on hydrogels of stiffness 10kPa and 250kpa, and plastic coated with collagen (P+C). Gene expression, of *p21* in (c) young and (d) senescent VSMCs when plated on hydrogels of stiffness 10kPa and 250kpa, and P+C. Gene expression of *IL6* in (e) young and (f) senescent VSMCs when plated on hydrogels of stiffness 10kPa and 250kpa, and P+C. Gene expression of *ACTA2* in (g) young and (h) senescent VSMCs when plated on hydrogels of stiffness 10kPa and 250kpa, and P+C. Gene expression of *TAGLN* in (i) young and (j) senescent VSMCs when plated on hydrogels of stiffness 10kPa and 250kpa, and P+C. N=7, One-way ANOVA with Tukey post hoc test, *p-value<0.05. Gene expression of (k) *ACTA2* and (l) *TAGLN* from young and senescent VSMCs plated on ECM synthesized from young VSMCS treated with EGFP control adenovirus (Young Ctrl), Sox9 overexpressing adenovirus (Sox9 OE), senescent VSMCs treated with shEGFP control adenovirus (Senescent Ctrl), and Sox9 knockout adenovirus (Sox9 KO). N=4, Unpaired Student's t-test. All gene expressions were quantified via RT-qPCR.

S. Figure 3

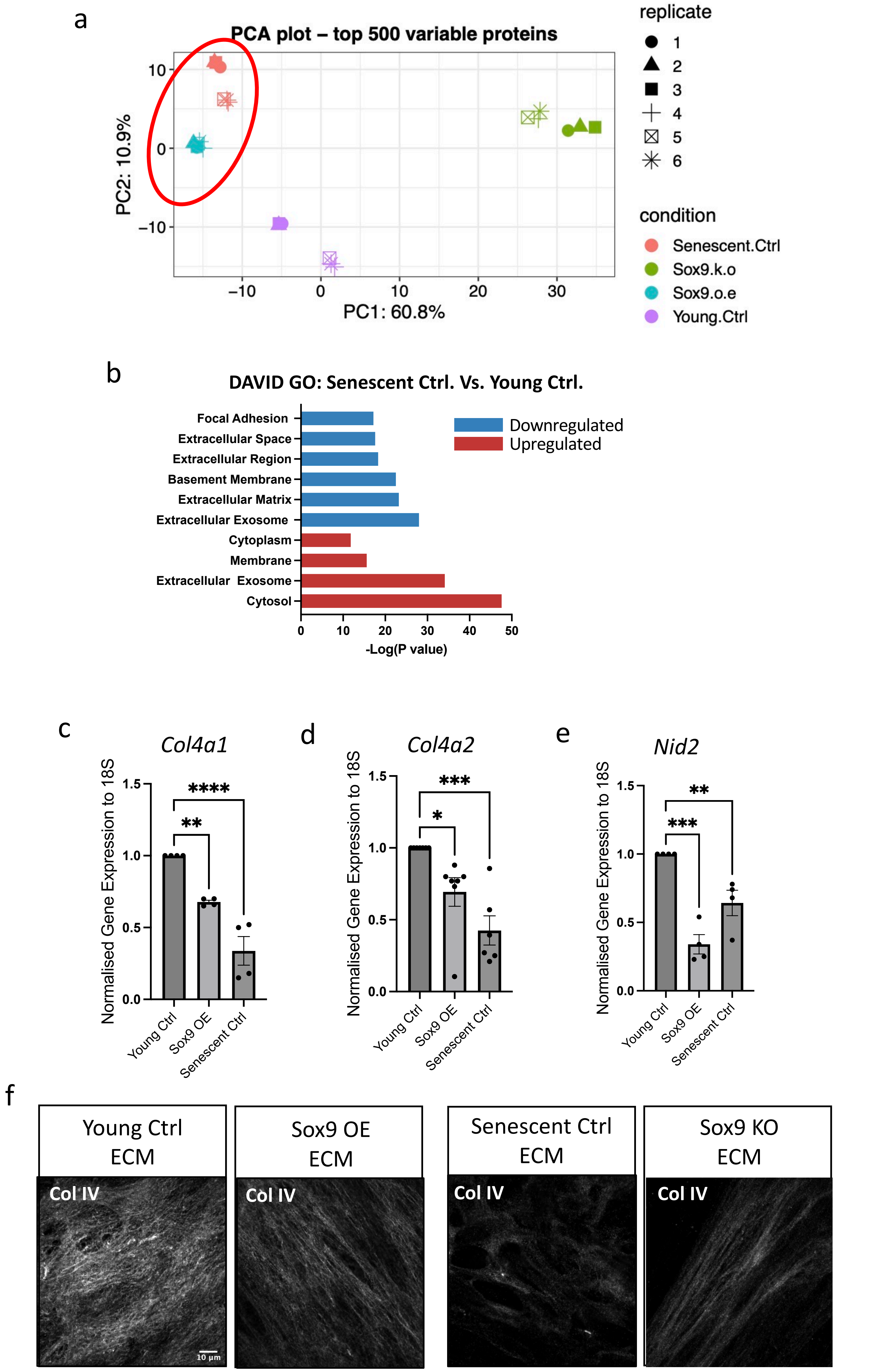


S Figure 4: (a) Principal component Analysis (PCA) plot of proteomics samples. Circle indicates clustered cell populations. (b) GO analysis showing pathways regulated by differentially expressed genes between young and senescent VSMCs. Gene expression, quantified via RT-qPCR of (c) Collagen (Col)4a1, (d), Col4a2, and (e) nidogen2 (Nid2) from young VSMCs transduced with EGFP control adenovirus (Young Ctrl), Sox9 overexpression adenovirus (Sox9 OE), and senescent cells transduced with shEGFP control adenovirus (Senescent Ctrl). N=4, One-way ANOVA with Tukey post hoc test, **p-value<0.01, ***p-value<0.005, p****<0.00001. (f) Immunofluorescence of Collagen 4 (Col IV) in ECM synthesized from Young Ctrl, Sox9 OE, Senescent Ctrl, and senescent VSMCs transduced with Sox9 knockout adenovirus (Sox9 KO).

S. Figure 4

a

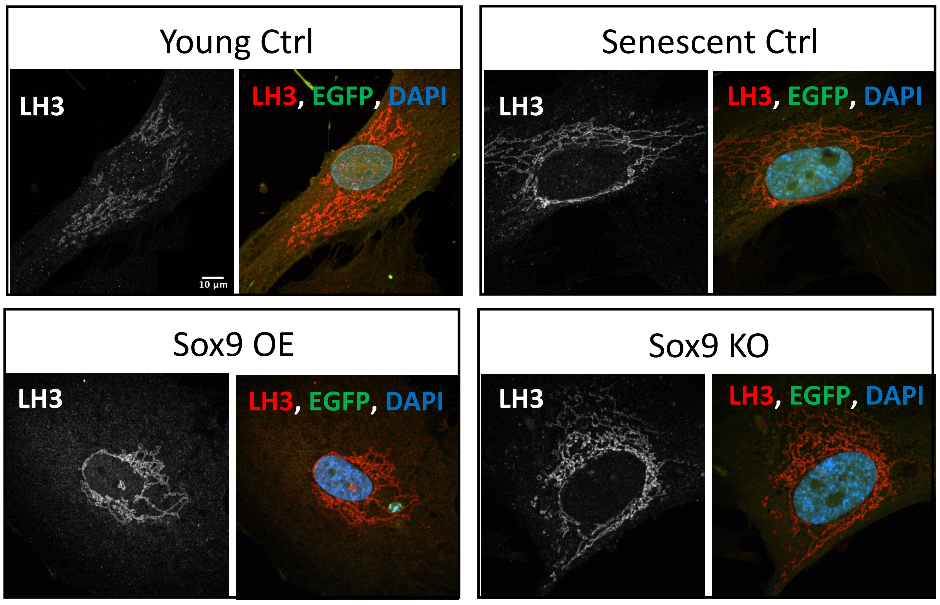


b

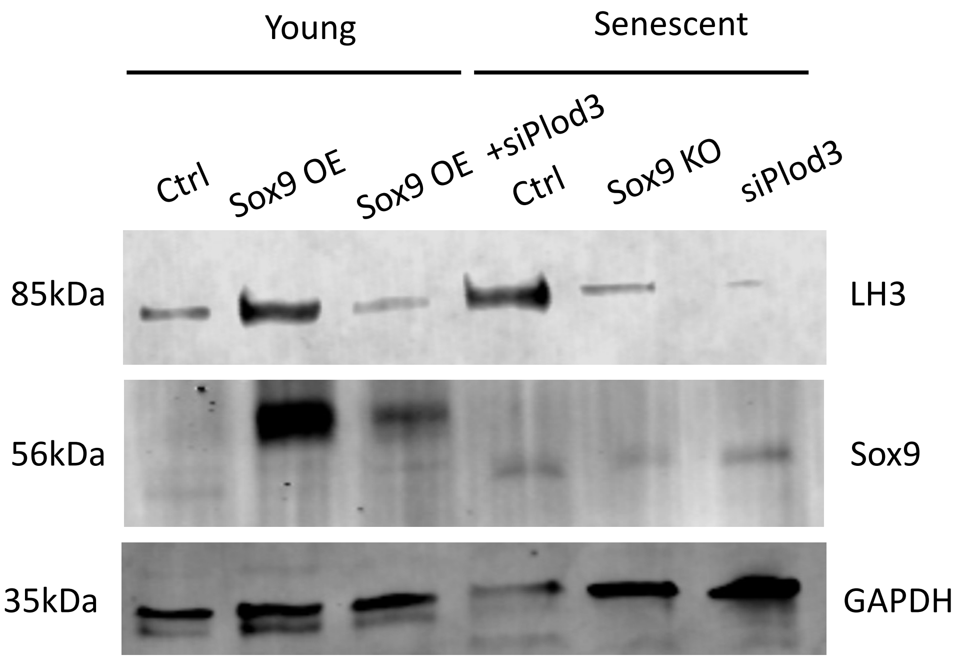


S Figure 5: (a) Representative immunofluorescence images of LH3 in young VSMCs transduced with EGFP control adenovirus (Young Ctrl), Sox9 overexpression (Sox9 OE), and senescent VSMCs transduced with shEGFP control adenovirus (Senescent Ctrl), and Sox9 knockout (Sox9 KO). LH3 is in grey and red, EGFP in green, and nuclear staining (DAPI) in blue. (b) Western blot depicting LH3 knockout (siPlod3) with Sox9 overexpression (Sox9 OE) and Sox9 knockout (KO) from protein lysate in young and senescent Vascular Smooth Muscle Cells.

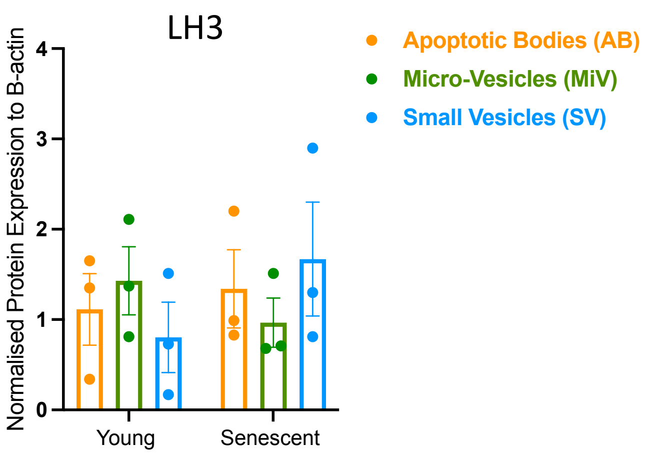


S. Figure 6: Protein quantification of LH3 from three types of extracellular vesicles in young and senescent Vascular Smooth muscle cells.
